# *PAX* fusion proteins deregulate gene networks controlling mitochondrial translation in pediatric rhabdomyosarcoma

**DOI:** 10.1101/2024.07.31.606039

**Authors:** Bhargab Kalita, Gerard Martinez-Cebrian, Justina McEvoy, Melody Allensworth, Michelle Knight, Alessandro Magli, Rita C R Perlingeiro, Michael A. Dyer, Elizabeth Stewart, Brian David Dynlacht

**Affiliations:** Department of Pathology and Perlmutter Cancer Institute, New York University School of Medicine, New York, NY, 10016, USA; Josep Carreras Leukaemia Research Institute, 08916, Badalona, Spain; Department of Developmental Neurobiology, St. Jude Children’s Research Hospital, 262 Danny Thomas Place, MS 323, Memphis, TN 38105-3678, USA; Department of Oncology, St Jude Children’s Research Hospital, Memphis, TN 38105-3678, USA; Department of Medicine, Lillehei Heart Institute, University of Minnesota, Minneapolis, MN, 55455, USA, Stem Cell Institute, University of Minnesota, Minneapolis, MN, 55455, USA; Genomic Medicine Unit - Sanofi, 225nd Ave Waltham, MA 02451

**Keywords:** Alveolar Rhabdomyosarcoma, Myogenic progenitors, TRMT5, MYCN, Mitochondrial metabolism, Roblitinib, Tigecycline, 3D/2D-adapted PDX models

## Abstract

Alveolar rhabdomyosarcoma (ARMS) patients harboring PAX3-FOXO1 and PAX7-FOXO1 fusion proteins exhibit a greater incidence of tumor relapse, metastasis, and poor survival outcome, thereby underscoring the urgent need to develop effective therapies to treat this subtype of childhood cancer. To uncover mechanisms that contribute to tumor initiation, we developed a novel muscle progenitor model and used epigenomic approaches to unravel genome re-wiring events mediated by PAX3/7 fusion proteins. Importantly, these regulatory mechanisms are conserved across established ARMS cell lines, primary tumors, and orthotopic-patient derived xenografts. Among the key targets of PAX3- and PAX7-fusion proteins, we identified a cohort of oncogenes, FGF receptors, and genes essential for mitochondrial metabolism and protein translation, which we successfully targeted in preclinical trials. Our data suggest an explanation for the relative paucity of recurring mutations in this tumor, provide a compelling list of actionable targets, and suggest promising new strategies to treat this tumor.

## Main

Alveolar rhabdomyosarcoma (ARMS) is a subtype of rhabdomyosarcoma, the most common pediatric soft tissue tumor (sarcoma). The disease is thought to arise from mesenchymal cells that fail to develop into skeletal muscle^1^. Compared to other subtypes, ARMS tumors are more aggressive and metastatic, leading to poorer prognoses and lower survival rates, and treatment options have not markedly improved over the past two decades. Nearly 80% of ARMS cases involve the fusion of the DNA binding domain of either of two master transcription factors (TFs), PAX3 and PAX7, with the activation domain of another transcription factor, FOXO1^2^. Expression of these fusion proteins subverts the normal myogenic development program, preventing differentiation of myogenic progenitors into mature skeletal muscle and leading to tumor formation^3^. Overall, patients with fusion protein (FP)-positive RMS exhibit a greater incidence of tumor relapse, metastasis, and poor survival outcomes^4, 5^ despite a lower mutational burden compared to another sub-type, embryonal RMS^6, 7^. While it is known that either fusion protein acts as an oncogene, it has not been possible to inhibit this protein through small molecules, and faithful, preclinical animal models for fusion-positive ARMS do not exist. Hence, developing experimental models that harness fusion protein expression within appropriate developmental windows will not only lead to a better understanding of genomic and epigenomic events mediated by the PAX-fusion proteins but also lead to effective therapies.

Previous reports have shown that PAX3-FOXO1 cooperates with other essential TFs, recruiting chromatin remodeling factors and histone-modifying enzymes to assemble super-enhancers (SE), which provide a platform for long-range interactions with promoters that drive expression of oncogenes in multiple cancers^3, 8^. However, mechanisms by which PAX7-FOXO1 abrogates normal myogenic differentiation and propels tumor initiation have not been explored in detail. Although previous reports have suggested that tumors with PAX3-FOXO1 are more aggressive and have poorer prognoses^9^, ARMS cases with PAX7 fusions are treated similarly^10^. We reasoned that understanding genome-wide changes in ARMS tumors bearing PAX7-FOXO1 could unravel key mechanistic differences. Further, while it is known that changes in tRNA synthesis, protein translation, and mitochondrial metabolism accompany oncogenic transformation in many tumors^11–17^, alterations in these processes have not be extensively studied in ARMS.

In an effort to explore how fusion proteins drive oncogenesis, we developed models in which both fusion proteins were expressed in muscle progenitors, allowing us to capture epigenetic changes and chromatin interactions and show how PAX fusion proteins re-organize chromatin architecture, in the absence of additional mutations, assembling super-enhancers that drive expression of key target genes. We identified key targets for both fusion proteins, novel regulatory elements for oncogenic drivers of ARMS, and actionable targets that include *FGFR4* and the mitochondrial translation machinery. Further, by generating robust PDX models from RMS patients, adapting them to two-dimensional growth, and performing preclinical testing in mouse models, we uncovered unanticipated, novel therapeutic strategies for high-risk ARMS.

## Results

### A novel model to study aberrant proliferation in ARMS tumors

To understand whether ARMS tumors resemble myogenic cells at a specific developmental stage, we compared normalized RNA-seq expression data from tumors with muscle cells undergoing differentiation. We compared data from human OCT4^+^ pluripotent stem cells, MSGN1^+^ pre-somitic cells, PAX7^+^ skeletal muscle progenitors, MYOG^+^ myoblasts, and multinucleated myotubes^18^ with transcriptomic data obtained from 26 patients with PAX fusion-positive ARMS^7^ and 4 orthotopic patient-derived xenografts^6^ (O-PDX) (Fig. 1A). Sample clustering using Pearson correlations, Euclidian sample distance, and principal component analysis (PCA) revealed a close relationship between gene expression profiles in ARMS tumors or O-PDXs and PAX7^+^ skeletal muscle progenitors (Figs. 1B, C, S1F). This finding is consistent with previous observations suggesting that ARMS tumor cells resemble cycling progenitors^19^ with expression signatures prevalent during the transition from embryonic to fetal muscle at 7–7.75 weeks of age^20^. We reasoned that expression of PAX3/7-FOXO1 fusion proteins in PAX7^+^ skeletal muscle progenitor cells might therefore capture key expression signatures in human ARMS without further introduction of additional oncogenic mutations.

**Figure 1.**
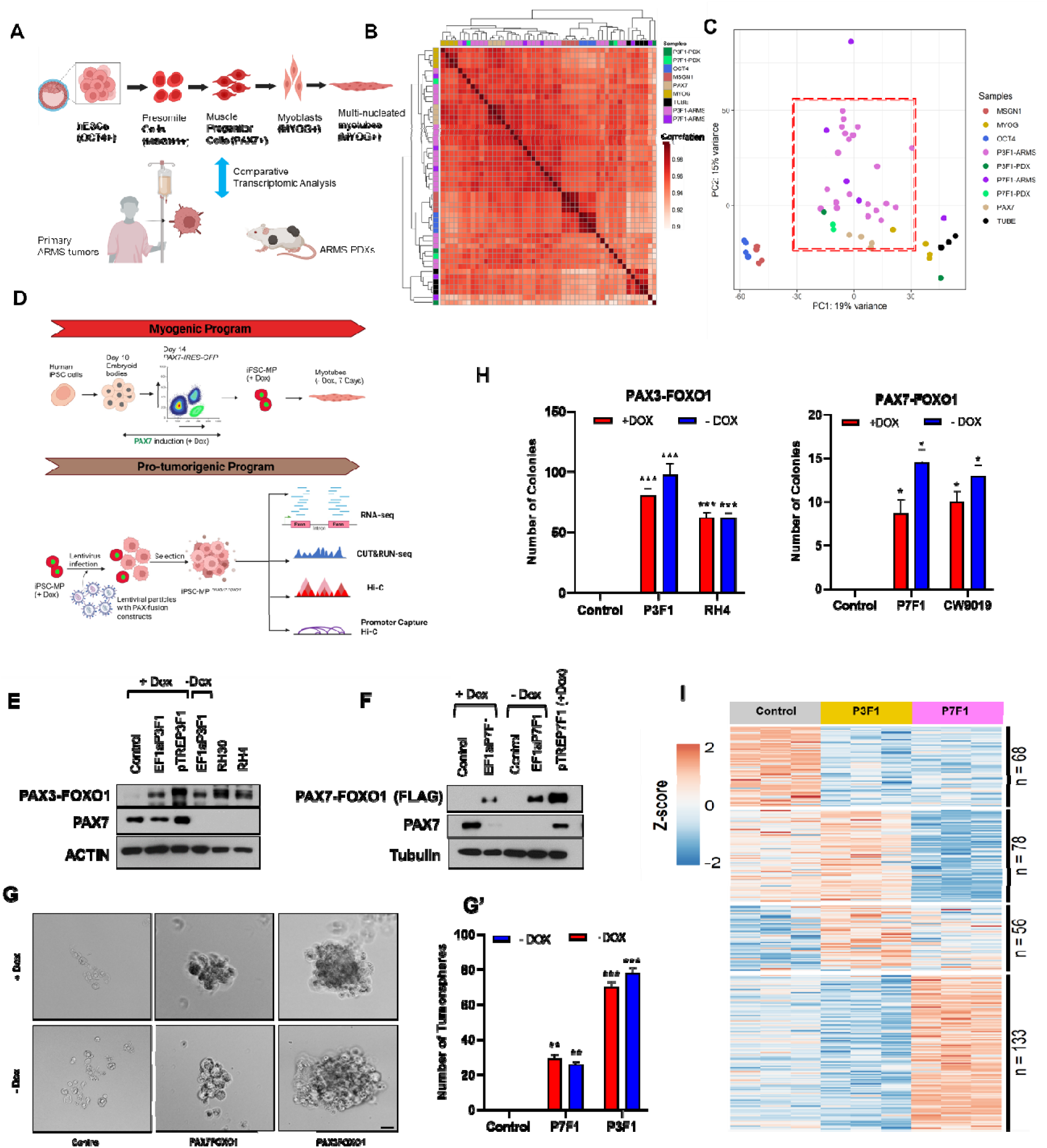
Development of iPSC models that express PAX3/7-FOXO1 fusion proteins. **A)** Scheme depicting comparative transcriptomic profiling of distinct muscle lineages reported previously^18^, primary ARMS tumors, and orthotopic patient-derived xenografts (O-PDX). **B)** Correlation Matrix plot comparing gene expression profiles of cells of different muscle lineage with alveolar rhabdomyosarcoma (ARMS) tumors and O-PDX. **C)** Principal component analysis plot showing distribution of RNA-seq normalized counts for OCT4+ pluripotent stem cells and distinct human myogenic stages, including MSGN1+ pre-somitic cells, PAX7+ skeletal muscle progenitor cells, MYOG+ myoblasts, and multinucleated myotubes with ARMS tumors and O-PDXs. **D)** Experimental strategy for generation of iPSC-MP ^PAX3/7FOXO1^ models and downstream characterization. **E-F)** Western blotting used to detect PAX3-FOXO1 (probed with PFM2.1 antibody), PAX7-FOXO1 (probed with FLAG antibody), and PAX7 in iPSC-MP ^PAX3/7FOXO1^ cells +/– Dox and in representative cell lines (RH30, RH4). **G)** Representative micrographs depicting tumor sphere formation in cells expressing PAX7-FOXO1 and PAX3-FOXO1 compared to fusion-negative control cells +/– Dox. Scale bar: 25 µm **G’)** Quantification of tumor sphere formation assay. n = 3 per group; error bar, SEM; **p < 0.01, ***p < 0.001, ANOVA, Dunnett’s test. **H)** Quantification of colony forming units in fusion-negative control cells, iPS-MP ^PAX3/7FOXO1^ cells (P3F1, P7F1), and indicated ARMS lines +/- Dox. n = 3 per group; error bar, SEM; *p<0.05, ***p < 0.001, ANOVA, Dunnett’s test. **I)** Heatmap depicting z-scores for muscle differentiation gene expression data from iPSC-MP ^PAX3/7FOXO1^ cells expressing PAX3-FOXO1 or PAX7-FOXO1 and controls. See also Figure S1.

We therefore utilized human induced pluripotent stem cells (iPSC) that differentiate into muscle progenitor-like cells (iPSC-MP) upon doxycycline (Dox) induction of PAX7 expression^21^. These cells are able to proliferate in the presence of inducer, but upon removal of Dox, iPSC-MP cells differentiate to form multi-nucleated myotubes^21, 22^ (Fig. 1D). We transduced these cells with lentiviruses encoding Flag epitope-tagged PAX3- or PAX7-FOXO1 fusion proteins (that precisely preserve fusion endpoints observed in tumors), allowing us to monitor their expression after selection and test the impact of their expression (Fig. 1D). Hereafter, we refer to this novel model as iPSC-MP^PAX3/7-FOXO1^. After removal of DOX, PAX7 expression ceases, and only the fusion protein is produced. Controls lacking the fusion protein were also established. We validated our iPSC-MP^PAX3/7-FOXO1^ models by detecting comparable levels of expression for each fusion protein in these cell lines and ARMS tumor cell lines expressing PAX3-FOXO1 (RH30, RH4) and PAX7-FOXO1 (CW9019) (Figs. 1E and S1A). PAX7-FOXO1 transcripts were produced at levels similar to those found in CW9019 (Fig. S1A), and using an antibody that specifically detects the PAX3-FOXO1 fusion protein, we showed that levels of this protein in our model cell line approximated those found in cognate ARMS cell lines (Fig. 1E). Since a PAX7-FOXO1-specific antibody was unavailable, we used antibodies against the epitope tag to confirm expression of this protein in our iPSC-MP^PAX7-FOXO1^ model (Fig. 1F).

Next, we investigated whether expression of either fusion protein is sufficient to drive abnormal growth and proliferation *in vitro*. Interestingly, we found that iPSC-MP^PAX3/7-FOXO1^ cells form tumor spheres *in vitro* when cultured under non-adherent conditions, suggesting they have acquired enhanced stemness and an ability to self-renew (Fig. 1G, G’). In contrast with parental controls, cells expressing the fusion proteins also readily displayed colony forming ability, a characteristic of ARMS cancer cell lines, including RH4 and CW9019 (Fig. 1H, S1B). Our data revealed that PAX3-FOXO1 expressing cells form larger numbers of tumor spheres and colony forming units (CFU) as compared to the PAX7-FOXO1 line, potentially signifying its enhanced aggressiveness. Taken together, our data indicate that expression of PAX3/PAX7 fusion proteins in myogenic precursors is able to model the aberrant growth and proliferation observed in ARMS tumor cells.

We next investigated whether fusion proteins enable tumor-associated gene expression signatures by performing transcriptome profiling of iPSC-MP^PAX3/7-FOXO1^ cells using RNA-seq. To more accurately test the role of fusion proteins, we performed our analysis in the absence of Dox (-3 days) to minimize interference from PAX7 in muscle progenitors. PCA analysis indicated strong concordance between replicates for populations expressing PAX3-FOXO1 or PAX7-FOXO1 and no overlap with muscle progenitor controls (Fig. S1C). The ability to cleanly separate each population suggests an underlying gene signature unique to each fusion protein. Our bulk RNA-seq analysis also revealed significant divergence from a normal muscle differentiation program (Fig. 1I), as well as down-regulation of genes associated with skeletal system development and the Wnt signaling pathway (Fig. S1D), a characteristic feature of rhabdomyosarcoma tumors^3, 23–25^. We found that expression of the PAX3-FOXO1 transgene alone led to up-regulation of 1783 genes and downregulation of 1451 genes compared to the control. Similarly, expression of PAX7-FOXO1 led to up-regulation of 1496 genes and down-regulation of 1869 genes (Fig. S1E, Table S1), indicating a pervasive role for both transcription factors in regulating gene expression. Consistent with our PCA analysis, cells expressing each fusion protein are marked by distinct gene expression programs, with 4500 genes differentially expressed between PAX3-FOXO1 and PAX7-FOXO1 cells (Fig. S1E’). Despite these differences, 488 genes are commonly up-regulated and 675 down-regulated in both models (Fig. 2A).

**Figure 2.**
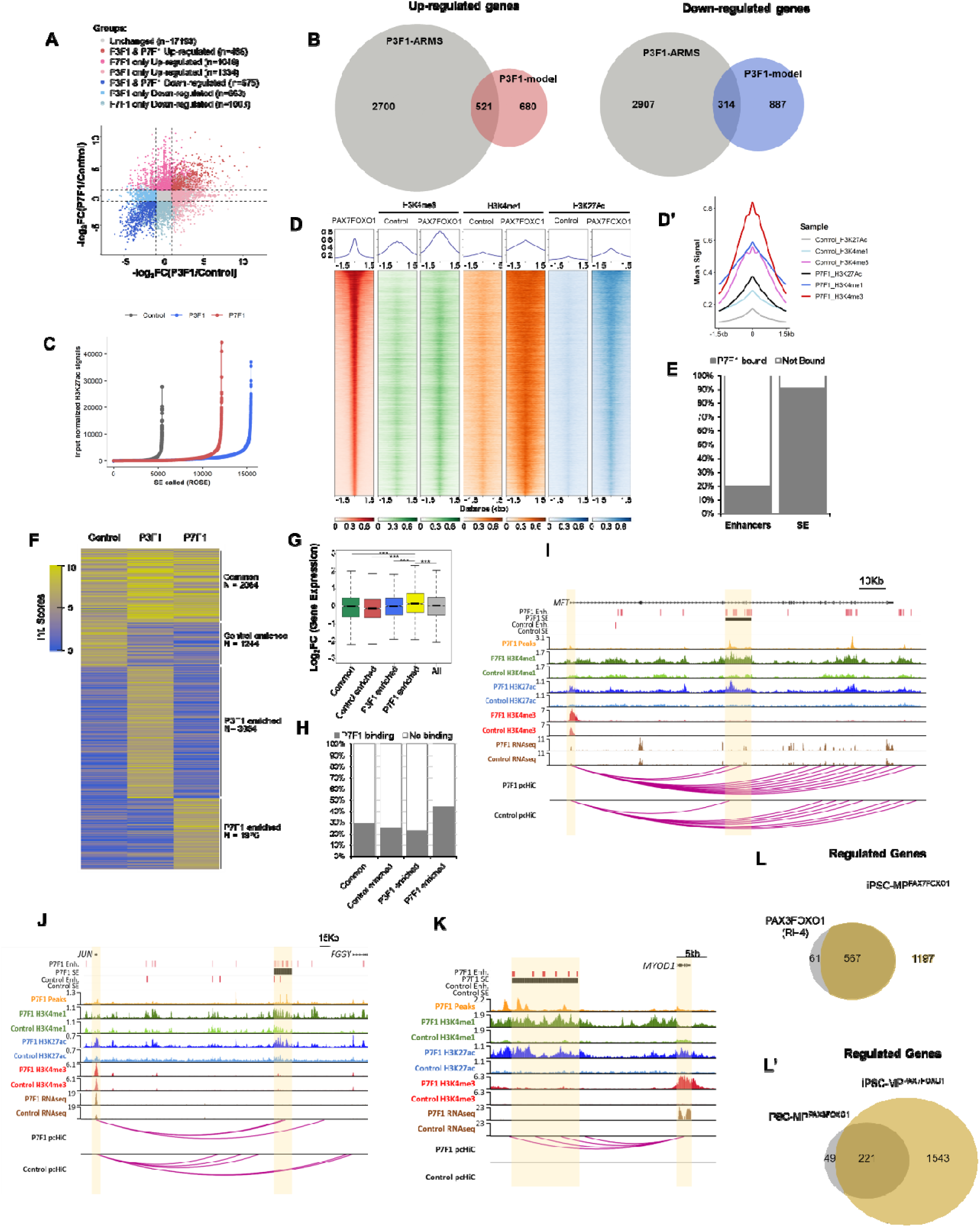
Genomic and epigenomic characterization of iPSC-MP^PAX3/7-FOXO1^ cells. **A** Plots for genes differentially expressed in iPSC-MP^PAX3-FOXO1^ and iPSC-MP^PAX7-FOXO1^ compared to fusion-negative control. **B)** Venn diagram depicting overlaps of transcriptomic data in PAX3-FOXO1-positive ARMS tumors and iPSC-MP^PAX3-FOXO1^ cells for down/up-regulated genes. **C)** Super enhancers were called for the indicated populations of iPSC-MP^PAX3/7-FOXO1^ and control cells using the ROSE algorithm. **D)** Composite genome-wide mean signal intensity plot for histone marks at PAX7-FOXO1 binding peaks. **D’)** Composite heatmap of enrichment for indicated histone modifications centered on PAX7-FOXO1 peaks (window indicates region between -1.5 and 1.5 kb relative to peak). **E)** Percentage of enhancers and super-enhancers (SE) bound by PAX7-FOXO1 transcription factor. **F)** *k*-means clustering of pCHiC promoter-enhancer interaction (Int) scores genes in iPSC-MP^PAX3/7-FOXO1^ cells. **G)** Box plot of normalized log_2_ fold-changes in gene expression of P-E interaction clusters common to all populations or enriched in fusion-negative controls, iPSC-MP^PAX3-FOXO1^, or iPSC-MP^PAX7-FOXO1^ cells. n = 8968 for all genes, n = 2064 for common, n = 1244 for control enriched, n = 3684 for P3F1 enriched, n = 1976 for P7F1 enriched; error bar, SEM; ***p < 10e^-17^, Welch’s t-test. **H)** Enrichment plots showing binding sites for PAX7-FOXO1 in the indicated enriched clusters. **I-K)** Genome browser tracks for regions encompassing **(I)** *MET*, **(J)** *JUN,* and **(K)** *MYOD1,* with promoter and super enhancers highlighted in yellow, showing CUT&RUN (H3K27Ac, H3K4me1, H3K4me3), RNA-seq, PAX7-FOXO1 binding sites, pCHiC probes, and high-confidence pCHiC interactions (magenta arcs) in control iPSC-MP and iPSC-MP^PAX7-FOXO1^ cells, after Dox removal. Super-enhancers are indicated for iPSC-MP^PAX7-FOXO1^ cells without Dox. Venn diagrams depicting overlap of genes regulated by PAX3-FOXO1 in RH4 **(L)** and iPSC-MP^PAX3-FOXO1^ **(L’)** or PAX7-FOXO1 mediated long-range P-E interactions. See also Figure S2.

We also examined whether expression changes instigated by PAX3/7-FOXO1 in iPSC-MP^PAX3/7-FOXO1^ cells overlap, or are distinct from, those found in ARMS tumors. Analysis of gene signatures in iPSC-MP^PAX7-FOXO1^ cells versus ARMS tumors expressing PAX7-FOXO1 revealed 400 of 1217 genes (33%) and 386 of 1611 genes (24%) to be commonly up-regulated and down-regulated, respectively, between the two groups (Fig. S1G), and a similar proportion was found to be commonly up-regulated and down-regulated in iPSC-MP^PAX3-FOXO1^ cells relative to ARMS tumors harboring PAX3-FOXO1 (43% up-regulated and 26% down-regulated)(Fig. 2B). Despite the expected differences between an *in vitro* cell line and patient-derived tissues, these overlapping gene sets are indicative of pathways strictly regulated by fusion proteins themselves, since additional mutations were not introduced in our iPSC-MP^PAX7-FOXO1^ cells. Commonly de-regulated genes include members of the HOX family, oncogenic drivers like FOXM1, FGFR4, and a key transcriptional regulator of mitochondrial metabolism, PPARGC1A (PGC-1α), each of which play critical roles in tumorigenesis and early development^26–29^. These findings validate our muscle progenitors as a novel, well-controlled model system useful for understanding the earliest steps of transformation associated with expression of the PAX3/7-FOXO1 fusion proteins.

### PAX fusion proteins alter chromatin architecture by assembling enhancers and super enhancers

To unravel fusion protein-associated chromatin regulatory mechanisms linked to tumorigenesis, we used Cut&Run to examine genome-wide binding by PAX3/7-FOXO1 and fusion protein-dependent changes in active histone marks (H3K27ac, H3K4me1 and H3K4me3) in our model. We defined 20,299 high-confidence binding sites for PAX7-FOXO1 and 1,122 binding sites for PAX3-FOXO1 in iPSC-MP^PAX3/7-FOXO1^ cells. The latter results indicated less pervasive binding than a previous study, which identified 5,129 PAX3-FOXO1 binding sites in RH4 cells^3, 23^, and therefore, we used these RH4 data to capture a broader compendium of binding sites. Using these data, we found that ∼2,874 genomic sites were commonly bound by both fusion proteins, whereas 17,425 and 2,255 sites were uniquely bound by PAX7-FOXO1 or PAX3-FOXO1, respectively (Fig. S2A). We found that both factors occupied promoter regions, distal intergenic, and intronic sites throughout the genome. Despite the existence of fewer binding sites around promoters, we found that cells expressing PAX3-FOXO1 exhibited a higher proportion of H3K4me3 deposition at promoter regions, as compared to controls and PAX7-FOXO1 expressing cells, potentially indicating the involvement of additional factors in gene activation in the former population (Fig. S2B).

We conservatively defined active enhancers by overlapping sites enriched for H3K27ac and H3K4me1 and removing active promoter regions with H3K4me3 enrichment. We identified 9,250 enhancers in iPSC-MP control cells, but this number expanded substantially in cells expressing PAX3-FOXO1 (36,243) and PAX7-FOXO1 (26,898). By implementing the ROSE algorithm^30^, we also defined super enhancer (SE) regions harboring the highest levels of H3K27ac deposition within the previously defined list of enhancer regions. Expression of either fusion protein led to a striking increase in super enhancer assembly compared to the control group, with PAX3-FOXO1 producing a considerably more robust response (Fig. 2C). Assembly of active enhancers and super enhancers is regulated in a cell-type and condition-specific manner, and, indeed, we found notable differences in the establishment of enhancers among these populations (Fig. S2C). Furthermore, we observed a significant enrichment in active enhancer marks (H3K27ac and H3K4me1) at PAX7-FOXO1 or PAX3-FOXO1 bound regions when comparing fusion protein expressing cells with controls (Figs. 2D, D’ and S2D, D’, D”). These results indicate that PAX7-FOXO1 can establish and activate enhancers *de novo* as previously observed for PAX3-FOXO1^3^, although the repertoire of enhancers targeted and activated by both proteins exhibited considerable divergence, again suggesting functional distinctions. Remarkably, we found that PAX7-FOXO1 occupied 20% and 90% of our annotated enhancers and super enhancers genome-wide, respectively, in iPSC-MP^PAX7-FOXO1^ (Fig. 2E).

To determine whether fusion with FOXO1 alters recruitment of PAX7 to chromatin, we compared sites bound by PAX7-FOXO1 with those bound by PAX7 in parental iPSC-MP cells^21^ used to generate our iPSC-MP^PAX7-FOXO1^ model. Interestingly, our analysis revealed that 3,376 genomic sites were commonly bound by both PAX7 and PAX7-FOXO1, whereas 31,523 and 16,923 genomic sites were uniquely bound by PAX7 and PAX7-FOXO1, respectively (Fig. S2E). When we focused on enhancers, we found that 834 sites were commonly occupied by both factors, whereas 482 and 4474 enhancer sites were uniquely bound by PAX7 and PAX7-FOXO1, respectively (Fig. S2E). Similarly, from a total of 511 super enhancers (SE) that we identified, 306 SE were commonly occupied by both factors, whereas 21 and 184 sites were uniquely occupied by PAX7 and PAX7-FOXO1, respectively (Fig. S2E). These results indicate that genome-wide differences between PAX7 and PAX7-FOXO1 binding are substantial, but the differences in SE binding may be more restricted. Motif discovery analysis revealed significant enrichment of canonical PAX7 binding motifs in FP-binding sites genome-wide (Fig. S2F).

Our previous studies demonstrated the ability of PAX7 to establish key regulatory nodes linking long-range promoter-enhancer interactions with gene expression during normal muscle differentiation^31^. We took advantage of an analogous approach to understand how fusion proteins effect genome-wide changes in chromatin architecture and gene expression in iPSC-MP^PAX3/7-FOXO1^ cells by performing HiC and promoter-capture HiC (pCHiC), which allowed us to map all genomic regions that interact with promoters. To enrich for functionally significant interactions, we restricted our analysis to include promoter interactions with active enhancers and super enhancers. We identified unique promoter-enhancer (P-E) interactions using *k*-means clustering of significant interactions (assessed using CHICAGO scores). From a total of 8,968 P-E interactions, 2,064 interactions were commonly observed in controls and in cells expressing PAX3-FOXO1 or PAX7-FOXO1. 3,684 and 1,976 P-E interactions were uniquely enriched in PAX3-FOXO1 and PAX7-FOXO1 expressing cells, respectively (Fig. 2F, Table S2).

Since comparatively little is known regarding the impact of PAX7-FOXO1 on gene expression, we focused on this factor by integrating our pCHiC and RNA-seq data. We found that PAX7-FOXO1 primarily regulates distant target genes through long-range promoter-enhancer contacts, frequently skipping over the nearest transcription start sites (TSS) (Fig. S3A). In cells expressing either fusion protein, we found that P-E interactions more strongly coincided with the occurrence of PAX7-FOXO1 binding events and were associated with increased gene expression (Fig. 2G, H). Importantly, binding of PAX7-FOXO1 at these regulatory regions, particularly super enhancers, also led to higher P-E interaction scores and associated target gene expression compared to controls (Fig. S3B-D), indicative of a pivotal role for the fusion protein in potentiating chromatin remodeling and strengthening promoter-enhancer interaction networks that enhance expression of its target genes.

### Fusion protein-mediated chromatin re-wiring enables expression of both myogenic differentiation and growth regulatory genes

Next, we examined whether PAX7-FOXO1 and PAX7 activate their respective target genes through common or distinct mechanisms. Since PAX7 and PAX7-FOXO1 bind to distinct enhancer and super enhancers regions (Fig. S2E), we reasoned that integrating promoter-enhancer (P-E) interaction networks could reveal genes that are differentially regulated by PAX7 and PAX7-FOXO1. Our data indicated that PAX7-FOXO1 occupied ∼50% of PAX7 bound enhancers to regulate expression of target genes via long-range PE interactions (Fig. S3E). Most interestingly, we noted that P-E interactions associated with PAX7-FOXO1 resulted in augmented expression of myogenic regulatory factors, including *MYOD1* and early lineage specification factors such as *SIX1*, which globally reprograms *MYOD1* to occupy genes that promote tumor growth instead of terminal muscle differentiation^32^. In fact, both TFs are highly expressed in RMS tumors^33, 34^. We further documented the global impact of PAX7-FOXO1 expression by demonstrating a role for the fusion protein in potentiating sustained and enhanced expression of a core group of target genes *after* PAX7 expression ceases (Fig. S3F).

ARMS tumors are characterized by a remarkable paucity of oncogenic mutations. Therefore, we linked our PAX3/7-FOXO1 binding and chromatin interaction data in an effort to understand its role in promoting transformation of muscle progenitors that lack known oncogenic mutations. Based on our analysis, we found that several oncogenes, including *MYCN*, *MET, JUN,* and *FOXM1* were bound and activated by PAX3-FOXO1 or PAX7-FOXO1 through specific P-E interactions (Figs. 3C, 2I, 2J and S3G). We also observed novel, PAX7-FOXO1-specific P-E interactions for several key muscle regulatory factors, MYOD1 and MYOG (Figs. 2K, S3H), and indeed, each of these genes was significantly up-regulated in these cells, corroborating observations in RMS. Importantly, a *MYOD1* enhancer cluster, located ∼25 kb upstream, was activated in cells harboring the PAX fusion protein and substantially augmented expression of the gene. This well-described “core” enhancer is activated in proliferating PAX7+ mouse muscle progenitors^31^ and is essential for *MYOD1* expression during early mouse development^35, 36^. Thus, the fusion protein establishes chromatin interactions to activate MYOD1, which maintains the cells in a highly proliferative state. We also found that a large cohort of muscle-specific genes was expressed in iPSC-MP^PAX3/7-FOXO1^ cells expressing either fusion protein, and these genes are known targets of both MYOD and MYOG^37–39^(Fig. 1I). Thus, both fusion proteins enhance MRF expression and potentiate a skeletal muscle-specific program observed in RMS, although a cohort of target genes were distinctly regulated by individual fusion proteins, indicating inherent differences between them (Fig. 2L, L’ Supplemental Table 6). Collectively, our analysis suggested how chromatin interactions instigated by PAX3-FOXO1 and PAX7-FOXO1 can simultaneously promote (1) maintenance of a proliferative progenitor state and aberrant growth in the absence of a high mutational burden and (2) expression of a muscle-specific program.

**Figure 3.**
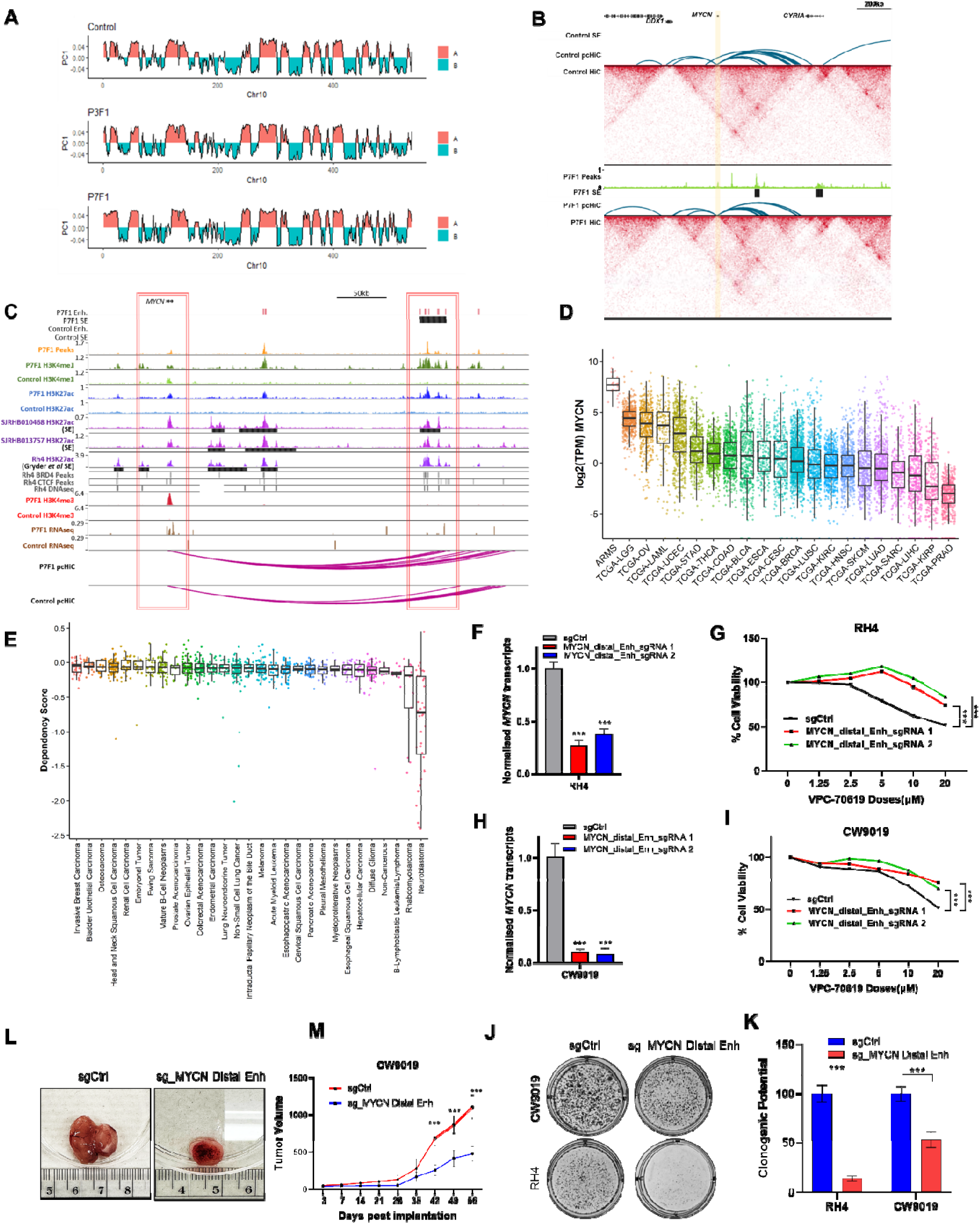
HiC and pCHiC analyses identify a distal enhancer downstream of *MYCN* essential for tumor growth. **A)** Compartment changes at chr10 for fusion negative control iPSC-MP vs. iPSC-MP^PAX3-FOXO1^ vs. iPSC-MP^PAX7-FOXO1^ cells. PC1 values (red: positive values; blue: negative values) indicating A and B compartments were generated at 250 kb resolution for comparison. H3K27ac signals were used to determine active chromatin associated with A compartments. **B)** HiC profile depicting topological-associated domains (TADs) encompassing *MYCN* gene (yellow bar) and high-confidence pCHiC interactions (blue arcs) in controls (iPSC-MP) and iPSC-MP^PAX7-FOXO1^ cells. PAX7-FOXO1 binding and super enhancers detected exclusively in iPS-MP^PAX7-FOXO1^ cells are indicated. **C)** Genome browser tracks encompassing *MYCN* locus (red, promoter), showing CUT&RUN (H3K27Ac, H3K4me1, H3K4me3), ChIP-seq (H3K27Ac, BRD4, CTCF in RH4), RNA-seq, DNase-seq (RH4), PAX7-FOXO1 binding sites, pCHiC probes, and high-confidence pCHiC interactions (magenta arcs) in control iPSC-MP and iPSC-MP^PAX7-FOXO1^ cells, after Dox removal. Super enhancers (SE; black bars) in iPSC-MP^PAX7-FOXO1^ cells without Dox, PAX7-FOXO1 positive O-PDX (SJRHB0104681, SJRHB013757)^6^, and RH4^3^ are shown. **D)** *MYCN* mRNA expression in TCGA data shows high levels of expression in RMS, among other tumor types. Abbreviations are as per TCGA (https://gdc.cancer.gov/resources-tcga-users/tcga-code-tables/tcga-study-abbreviations). **E)** Box plots showing average dependency scores for MYCN. **F)** *MYCN* transcript levels in two independent CRISPRi clones compared to sgCtrl in RH4 cells. n = 3 per group; error bar, SEM; ***p < 0.001, ANOVA, Dunnett’s test. **G**) Cell viability curves for RH4 control (sgCtrl) and two CRISPRi clones exposed to indicated doses of the MYCN inhibitor VPC-70619. n = 3 per group; error bar, SEM; ***p < 0.001, ANOVA, Dunnett’s test. **H**) *MYCN* transcript levels in two independent CRISPRi clones compared to sgCtrl in CW9019 cells. n = 3 per group; error bar, SEM; ***p < 0.001, ANOVA, Dunnett’s test. **I)** Cell Viability curve of Cas9 Ctrl and two CRISPRi clones of CW9019 cells treated with indicated doses of the MYCN inhibitor VPC-70619. n = 3 per group; error bar, SEM; ***p < 0.001, ANOVA, Dunnett’s test. **J)** Colony assays comparing clonogenic growth potential in sgCtrl and CRISPRi-silenced cell populations of CW9019 and RH4. **K)** Quantification of results in panel J. n = 3 per group; error bar, SEM; ***p < 0.001, ANOVA, Dunnett’s test. **L)** Representative tumors from sgCtrl and CRISPRi-silenced CW9019 cell lines. **M)** Tumor growth curves for CW9019 clones stably expressing control sgRNAs and sgRNAs specific to *MYCN* SE. n = 3 mice per group; error bar, SEM; ***p < 0.001, ANOVA, Sidak test. See also Figure S3.

### Fusion protein-mediated re-wiring of *MYCN* locus is essential for tumorigenesis

We also performed HiC to understand if PAX3 and PAX7 fusion proteins alter properties of higher order genome organization, including chromatin compartments and topologically associated domains (TADs). Based on our HiC analysis, we noted that a total of ∼15% of compartments switched from A to B (silencing) and from B to A (activation) in cells expressing the fusion proteins (Fig. 3A, S3I). This observation is in line with previous reports^40–42^ which show that compartments/TADs are largely conserved across cell types. In addition, we found approximately 2000 TADs to be associated with each of our analyzed samples (Table S3). We used TADcompare to identify differences in TAD boundaries across cell types (Fig. S3J). In some cases, altered TAD boundaries were associated with key oncogenic drivers of RMS like *MYCN*, where insulation of a sub-TAD was observed following occupancy of the fusion protein at a distant super-enhancer (Fig. 3B). Inspection of pCHiC interactions within this TAD revealed a novel regulatory element approximately ∼200kb downstream of the *MYCN* promoter in cells expressing PAX7-FOXO1 (Fig. 3C). Interestingly, this super enhancer recruited PAX7-FOXO1 and contacted the *MYCN* promoter through long-range PE interactions, which were lacking in the control (Fig. 3C). The functional relevance of this novel super enhancer had not been investigated previously, although SEs ∼90 kb and ∼56 kb downstream of *MYCN* have been defined for PAX3-fusion positive ARMS^3^ and neuroblastoma^43^, respectively. Interestingly, we found that BRD4 and CTCF are recruited in RH4 cells at sites that coincide with PAX7-FOXO1 peaks in our model, and acquisition of H3K27Ac marks and open chromatin states in FP-positive tumors and PDX models in this particular genomic region further substantiates its regulatory and clinical importance in ARMS. The presence of these regulatory elements was associated with increased *MYCN* expression, which was elevated in ARMS compared to other aggressive cancers (Fig. 3D). Moreover, DepMap analysis revealed a low dependency score for *MYCN* in RMS cells compared to most other cancers (Fig. 3E), indicating an essential role for MYCN in RMS tumor survival and progression.

To understand the biological role of the SE element ∼200kb downstream of the *MYCN* gene and its essentiality in RMS, we used CRISPRi to silence this enhancer region in two representative ARMS cell lines, RH4 and CW9019. Stable expression of two distinct sequence-specific guide RNAs and the Cas9-KRAB-MeCP2 repressor resulted in silencing of *MYCN* expression in both ARMS cell lines (Fig. 3F, H). Further, stable silencing led to dose-dependent resistance to VPC-70619, a MYCN inhibitor, as compared to controls, confirming both the specificity of our silencing strategy and the dependency of ARMS cells on *MYCN* expression (Fig. 3G, I). Next, we tested the impact of MYCN silencing on growth of these tumor cell lines using adhesion-dependent growth assays. We observed significant reductions in clonogenic growth of CRISPRi silenced ARMS cells, as compared to their corresponding controls (Fig. 3J, K). This suggests a critical role for the MYCN super enhancer in regulating the self-renewing capabilities of ARMS cells. To further test the functional role of the enhancer region in tumorigenicity, we subcutaneously xenografted control and CRISPRi silenced cells in NOD/SCID mice and scored for tumor growth. Our data confirmed the role of these enhancer elements in ARMS tumorigenicity, as CRISPRi silencing significantly delayed tumor growth, and implanted cells formed smaller tumors compared to control groups after implantation (Fig. 3L, M). Collectively, our data suggest a functional link between fusion protein-mediated assembly of specific enhancer and super-enhancer clusters and expression of key oncogenic drivers, such as MYCN, in ARMS. Our data may also explain why *MYCN* is amplified in a subset, but not a majority, of ARMS tumors^7^, since fusion-protein mediated enhancement of expression may obviate the need for gene amplification.

### RMS tumor display high mitochondrial OXPHOS activity

Our foregoing studies using iPSC-MP^PAX3/7-FOXO1^ cells unveiled key, unexplored FP targets and novel regulatory elements activated in a fusion protein-dependent manner, and we further leveraged these data in an effort to identify druggable pathways activated in ARMS. Remarkably, upon clustering differentially expressed genes in these models using Gene Ontology (GO), we observed striking deregulation of a large cohort of mitochondrial genes as a consequence of fusion protein expression (Fig. S4A). From a total of 1046 expressed mitochondria-related genes in MitoCarta3.0, 305 genes and 259 genes were uniquely up-regulated in cells expressing PAX7-FOXO1 and PAX3-FOXO1, respectively, and 203 genes were found to be commonly up-regulated (Fig. S4A). Accordingly, we observed higher basal and maximal oxygen consumption rates (OCR) in our models expressing the fusion proteins compared to fusion-negative controls (Fig. S4B, B’). These data suggest that expression of either fusion protein is sufficient to drive aberrant OXPHOS phenotypes in RMS. Importantly, when we integrated our factor binding and pCHiC datasets, we showed that these fusion TFs directly target and activate mitochondrial metabolism genes (∼266 genes for PAX7-FOXO1; ∼63 genes for PAX3-FOXO1), (Tables S2, S4, S6).

Our analyses prompted us to further dissect the functional role of mitochondria in ARMS patients by comparing transcriptomic profiles of mitochondria-related genes derived from a panel of four O-PDXs and normal human skeletal muscle myoblasts (HSMM). We observed up-regulation of 793 mitochondrial genes in the fusion-positive PDXs, whereas 249 genes exhibited downregulation (Fig. 4A). GO terms associated with these genes included mitochondrial translation and respiratory electron transport chain assembly. Further, gene-set enrichment analysis (GSEA) indicated enrichment of an overlapping group of mitochondria-related genes in O-PDXs samples (Fig. 4B). Remarkably, we also observed PAX3-FOXO1 binding to the enhancer of *PPARGC1A*, a master regulator of mitochondrial metabolism, together with P-E interactions and a change in TAD boundaries (Fig. 4C, Table S3). *PPARGC1A* is a transcriptional coactivator that regulates a wide array of genes involved in OX-PHOS metabolism as well as mitochondrial biogenesis and function,^44^ and its upregulation has been linked to therapy resistance in multiple malignancies^45, 46^. However, little is known about its role in rhabdomyosarcoma. We found that *PPARGC1A* was significantly up-regulated in ARMS tumors and O-PDXs samples, as compared to most cancers in the TCGA database (Fig. 4C’). Further, its key transcriptional targets, *NRF1*, *TFAM*, *TFB1M,* and cytochrome c oxidase subunits, responsible for initiating mitochondrial biogenesis and function^47, 48^, were also found to be up-regulated in transcriptomic datasets derived from ARMS patient tumors^7^ and O-PDX models^6^ (Table S5). GSEA analysis indicated that highly over-represented target gene clusters involve mitochondrial translation and include mitochondrial ribosomal subunits and leucyl-tRNA synthetase (*LARS2*) (Table S4).

**Figure 4.**
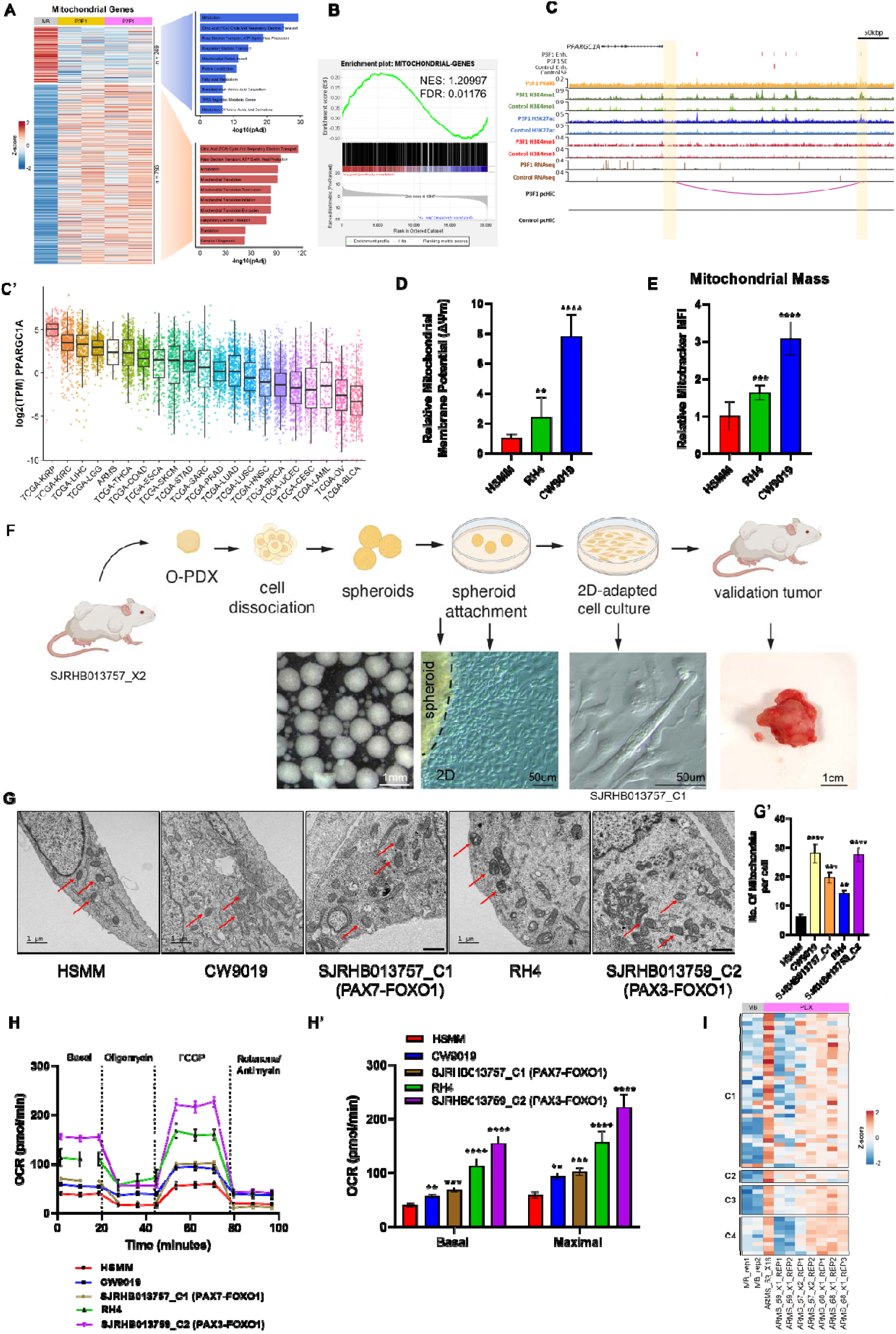
Investigating mitochondrial alterations in ARMS. **A)** Heatmap showing transcriptomic profile of mitochondria-related genes in human myoblasts (MB) from healthy donor, PAX3-FOXO1 (P3F1) and PAX7-FOXO1 (P7F1) O-PDX samples. **B)** Gene-set enrichment analysis (GSEA) of transcriptomic data from PAX-fusion O-PDX showing normalized enrichment score (NES) of selected gene ontology terms (mitochondria-associated genes). **C)** Genome browser tracks encompassing *PPARGC1A* locus (yellow, promoter and distal enhancer), showing CUT&RUN (H3K27Ac, H3K4me1, H3K4me3), RNA-seq, PAX3-FOXO1 binding sites, pCHiC probes, and high-confidence pCHiC interactions (magenta arcs) in control iPS-MP and iPS-MP^PAX3-FOXO1^ cells after Dox removal. **C’)** *PPARGC1A* mRNA expression in TCGA shows high level of expression in RMS and other tumor types. TCGA abbreviations shown in Fig. 3D. **D)** Quantification of TMRM dye uptake in HSMM, RH4, and CW9019 cells. n = 3 per group; error bar, SEM; **p < 0.01, ****p < 0.0001, ANOVA, Dunnett’s test. **E)** Quantification of mitochondrial mass using Mitotracker dye uptake in HSMM, RH4, and CW9019 cells. n = 3 per group; error bar, SEM; ***p < 0.001, ****p < 0.0001, ANOVA, Dunnett’s test. **F)** Representative workflow showing generation of 3D spheroids and 2D-adapted PDX models from SJRHB013757_X2 (PAX7-fusion positive patient’s O-PDX) and its validation. **G)** TEM micrographs showing mitochondria (red arrows) in HSMM, CW9019, SJRHB013757_C1 (PAX7-FOXO1), RH4, and SJRHB013759_C2 (PAX3-FOXO1) cells. Scalebar: 1μM G**’)** Quantification of mitochondria in HSMM, CW9019, SJRHB013757_C1, RH4, and SJRHB013759_C2 cells. error bar, SEM; **p < 0.01, ***p < 0.001, ****p < 0.0001, ANOVA, Dunnett’s test **H)** Seahorse Assay profiles for HSMM, CW9019, SJRHB013757_C1 (PAX7-FOXO1), RH4, and SJRHB013759_C2 (PAX3-FOXO1) cells, as indicated, using Cell Mito Stress assay. **H’)** Quantification of basal and maximal oxygen consumption rates (OCR) shown in panel G. n = 3 per group; error bar, SEM; **p < 0.01, ***p < 0.001, ****p < 0.0001, ANOVA, Dunnett’s test **I)** Heatmap showing protein levels of mitochondrial respiratory complex proteins in human myoblasts, PAX3-FOXO1 (59_X1, 63_X16) and PAX7-FOXO1 (57_X2, 68_X1) O-PDX samples. See also Figure S4.

To further explore the role of mitochondrial function in ARMS, we measured differences in mitochondrial membrane potential in two representative ARMS cell lines, RH4 (PAX3-FOXO1) and CW9019 (PAX7-FOXO1), by assessing tetramethylrhodamine methyl ester (TMRM) dye uptake. We found that ARMS cells possessed a higher potential difference across mitochondrial membranes, as compared to human myoblasts, to generate ATP needed to meet the greater energy demands in tumor cells (Fig. 4D). Further, Mitotracker staining and FACS analysis indicated a significantly higher mitochondrial mass in ARMS cells compared with normal human myoblasts (Fig. 4E). In agreement, transmission electron microscope (TEM) analysis also revealed significant increases in the number of mitochondria in cells harboring PAX fusion proteins compared to normal controls (Fig. 4G, G’). To further examine a role for mitochondrial respiratory complexes, we investigated mitochondrial respiration in ARMS cell lines and normal human myoblasts using Seahorse assays. We also developed and utilized a novel set of 2D-adapted PDX models (SJRHB013757_C1 and SJRHB013759_C2) with analogous tumor heterogeneity and genetic composition as found in their matched PDX tumor models and ARMS patients^49^ (Fig. 4F). An in-depth profiling of 3D/2D-adapted PDX models derived from other solid tumors has been submitted as a separate study (McEvoy et al., unpublished). Our data indicated that higher basal and maximal oxygen consumption rates (OCR) were associated with ARMS tumor cells and PDX models compared to normal, human myoblasts (Fig. 4H, H’). Addition of mitochondrial complex inhibitors and decouplers demonstrated the enhanced ability of fusion-positive ARMS and 2D-adapted PDX models to respond to mitochondrial stress with an increased OCR rate compared to normal myoblasts (Fig. 4H). Strikingly, when we examined proteome-wide expression data derived from each of these O-PDX models^6^, we observed that the augmentation of basal and maximal OCR rates in ARMS and 2D-adapted PDX cells coincided with the increased abundance of proteins associated with mitochondrial complexes I-V, in agreement with their altered gene expression (Fig. 4A, B, I, Table S5).

To delineate the contribution of MYCN to the observed increases in OXPHOS metabolism in ARMS cells, we depleted MYCN but found that there was no impact on OCR rates (Fig S4C, C’, D, D’). Taken together, we conclude that ARMS tumors rely on increases in OXPHOS activity to compensate for altered energy requirements, and these metabolic changes reflect the impact of fusion protein-mediated enhancement of genes encoding mitochondrial translation and OXPHOS machinery.

### Mitochondrial tRNAs are up-regulated in ARMS tumors

Changes in protein translation accompany metabolic demands during tumorigenic initiation and progression, and the cell accommodates such changes through increased expression of tRNAs, with codon biases for specific amino acids leading to perturbation of mitochondrial metabolism in many malignancies^11–15, 50^. We reasoned that tRNA availability and/or the abundance of components of the mitochondrial translation machinery could represent a therapeutic vulnerability for certain tumors whose growth is constrained by their metabolic demands.

We therefore investigated the mechanisms connecting fusion protein production with aberrant expression of mitochondrial proteins in ARMS. Increased expression of specific tRNAs has been noted in human cancers, with altered codon usage matched to the cognate tRNA species^51^. Since our data suggested that mitochondrial protein expression was up-regulated in tumors, we analyzed the expression profiles of 22 mitochondrial tRNAs using transcriptomic data from a panel of 26 patients with ARMS tumors^7^. Remarkably, our data indicated that mitochondrial tRNAs are frequently upregulated in both ARMS tumor types and iPSC-MP^PAX3/7-FOXO1^ cells (Fig. 5A, A’). We therefore selected a subset of tRNAs most highly up-regulated in ARMS and validated our observations by performing mitochondrial tRNA-specific qPCR in a panel of ARMS O-PDX samples and normal myoblasts after removal of base modifications. Our data indicated significant up-regulation of specific mitochondrial leucine tRNA isoacceptors (*TRNL1*, *TRNL2*) and a glycine tRNA (*TRNG*) in O-PDX samples when compared to human myoblasts (Fig. 5B). Using the R package, coRdon, we analyzed mitochondrial respiratory complex genes (from THE HUGO database) and observed significant enrichment in the frequency of Leu codons comprising the consensus ‘CUN’ that are cognate to the TRNL2 anticodon in genes up-regulated in O-PDX and ARMS compared to mitochondrial complex genes that do not vary in expression upon fusion protein expression (Fig. S5A, A’). This finding suggests that ARMS tumors preferentially express mitochondrial tRNAs that promote biased usage of specific leucine codons to enhance expression of mitochondrially encoded genes.

**Figure 5.**
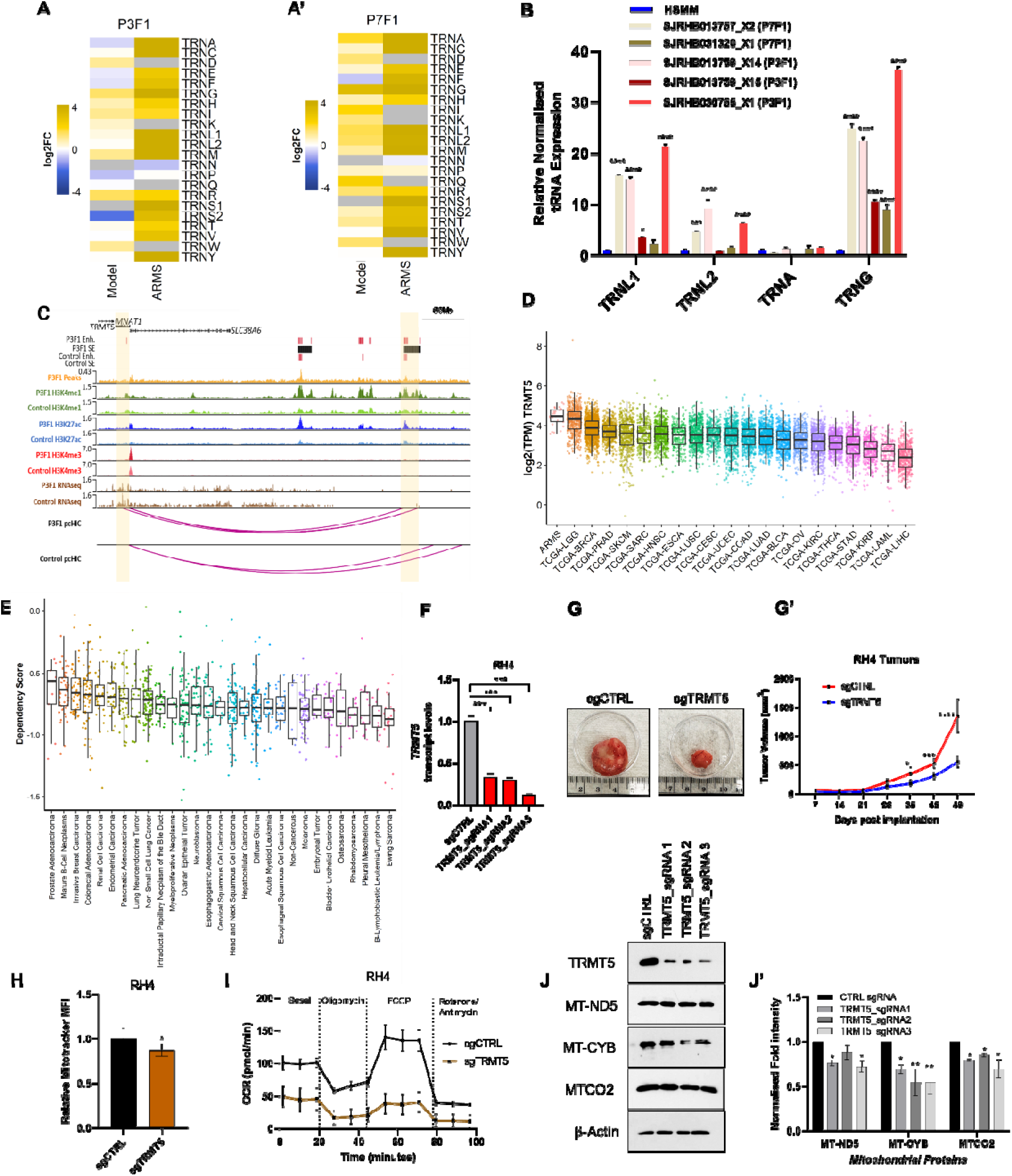
*TRMT5* plays an essential role in ARMS growth. **A, A’)** Heatmap of expression profiles of 22 mitochondrial tRNAs in the indicated iPSC-MP^PAX3/7-FOXO1^ populations (model) and corresponding primary tumors normalized to human myoblasts. **B)** Quantitative qPCR data for mitochondria-specific tRNA expression in human myoblasts and PAX-fusion driven O-PDX samples. SJRHB013759_X14 and SJRHB013759_X15 are O-PDX samples, collected from two different metastatic sites i.e. chest and omentum respectively, from the same patient. Details can be found in https://cstn.stjude.cloud/search/. n = 3 per group; error bar, SEM; *p < 0.05, ***p < 0.001, ****p < 0.0001, ANOVA, Dunnett’s test. **C)** Genome browser tracks encompassing *TRMT5* locus (yellow, promoter), showing CUT&RUN (H3K27Ac, H3K4me1, H3K4me3), RNA-seq, PAX3-FOXO1 binding sites, pCHiC probes, and high-confidence pCHiC interactions (magenta arcs) in control iPS-MP and iPS-MP^PAX3-FOXO1^ cells after Dox removal. Super-enhancers detected in iPS-MP^PAX3-FOXO1^ cells without Dox are indicated. **D)** TCGA data showing high levels of *TRMT5* expression in ARMS. Abbreviations are given in Fig. 3D. **E)** Average box plots of dependency scores for *TRMT5* in RMS indicating greater dependency on *TRMT5* for survival. **F)** Transcript levels of *TRMT5* following CRISPRi silencing in RH4 cells using three independent sgRNAs targeting the promoter region. n = 3 per group; error bar, SEM; ***p < 0.001, ANOVA, Dunnett’s test. **G)** At terminal end-points, representative images depicting RH4 tumors stably expressing control and TRMT5-specific gRNAs. **G’)** Tumor growth curve of RH4 stably expressing control and sgRNAs specific to TRMT5. n = 3 mice per group; error bar, SEM; *p < 0.05, ***p < 0.001, ****p < 0.0001, ANOVA, Sidak test. **H)** Quantification of mitochondrial mass using Mitotracker dye uptake in RH4 cells stably expressing control and TRMT5-specific gRNAs. n = 3 per group; error bar, SEM; *p < 0.05, Student’s t test. **I)** Seahorse assays showing oxygen consumption rates (OCR) for RH4 cells stably expressing control and TRMT5-specific sgRNAs using Cell Mito Stress kit. **J)** Western blot images of TRMT5, MT-ND5, MT-CYB, MTCO2 and β-Actin in RH4 clones expressing control and TRMT5-specific sgRNAs. **J’)** Quantification of data in panel J. n = 3 per group; error bar, SEM; *p < 0.05, **p < 0.01, ANOVA, Dunnett’s test. See also Figure S5.

### *TRMT5* is a novel target of fusion proteins driving ARMS tumorigenesis

During the process of maturation, tRNAs undergo several post-transcriptional modifications, and we found that several genes encoding modifying enzymes, including *TRMT5*, were up-regulated in iPSC-MP^PAX3/7-FOXO1^, ARMS tumors, and/or O-PDXs (Table S5). TRMT5, a methyltransferase, localizes to mitochondria, where it promotes N1-methylation of the guanosine residue at position 37 (G37) of mitochondrial tRNA^Leu^ ^(CUN)^ (*TRNL2*)^52^, which affects translational fidelity, ribosome stalling, and codon usage bias, and is essential for viability^53–55^. Most studies on TRMT5 have been performed in bacteria or yeast. It is not known whether TRMT5-mediated methylation alters the levels of target tRNAs or how it impacts proliferation and oncogenesis, and its role has not been investigated in RMS or other cancers. We found that *TRMT5* is a target of both oncogenic fusion proteins, and our analysis revealed super enhancers bound by PAX3-FOXO1 and PAX7-FOXO1 downstream of *TRMT5* locus, which exhibited long-range interactions with the *TRMT5* promoter (Fig. 5C, S5B, Table S3). Gene expression analyses utilizing the TCGA repository revealed that TRMT5 is highly expressed in RMS tumors as compared to other cancer types (Fig. 5D). Additionally, DepMap-based cancer dependency analysis indicated that rhabdomyosarcomas have a significantly lower dependency score compared to other tumors, indicating a greater dependency on *TRMT5* for survival (Fig. 5E). To validate these observations, we performed western blot analysis of *TRMT5* expression in a panel of ARMS cell lines, including CW9019, RH4, RH30, and two 2D-adapted PDX models expressing PAX3-FOXO1 and PAX7-FOXO1. Notably, this analysis indicated that TRMT5 is upregulated in all tumors and 2D-adapted PDX models expressing either fusion protein as compared to normal myoblasts, and *TRMT5* is more highly expressed in PAX3-FOXO1 positive tumor cells (Fig. S5C). This observation is consistent with our proteomics data from O-PDX samples (Fig. S6A’).

To understand the functional role of *TRMT5* in ARMS pathogenesis, we performed CRISPRi-based silencing using sequence-specific sgRNAs targeting the *TRMT5* promoter in RH4 cells. We tested three different sgRNAs, each of which potently suppressed *TRMT5* transcription and reduced protein levels in ARMS cells (Fig. 5F, J, J’), and examined growth using colony forming assays. Our data indicated that *TRMT5* silencing significantly reduced the clonogenic potential of RH4 cells over a range of cell densities (Figs. S5D, D’). This suggests a critical role for TRMT5 in regulating the self-renewal capabilities of ARMS cells. To confirm the functional role of TRMT5 in ARMS tumorigenicity, we subcutaneously implanted control and TRMT5-silenced cells in NOD/SCID mice and assessed the tumorigenicity of transplanted cells. Silencing of TRMT5 significantly delayed tumor growth, resulting in substantially smaller tumors (∼3-fold reduction in volume) as compared to controls (Fig. 5G, G’). To gain mechanistic insights into the role of TMRT5 in mitochondrial function, we quantified Mitotracker fluorescence with flow cytometry in cells expressing control sgRNAs and TRMT5 targeted sgRNAs. We found that TRMT5 silencing resulted in significant reductions in mitochondrial mass and corresponding, dramatic decreases in basal and maximal OCR rates as compared to controls, reflecting severe reductions in the process of oxidative phosphorylation (Fig. 5H, I).

TRMT5 is a tRNA methyltransferase, and therefore, we sought to link the levels of this enzyme with one of its targets, mitochondrial *TRNL2*. Interestingly, TRMT5 silencing led to a significant reduction in TRNL2 tRNA expression in ARMS cells, resulting in decreased levels of mitochondrial OXPHOS proteins, MT-ND5, MT-CYB, and MTCO2 (Figs. S5E, 5J, J’). Since TRNL2 encodes the mitochondrial tRNA^Leu^ ^(CUN)^ isoacceptor, this result provides a plausible explanation for the enhanced expression of mitochondrial respiratory complex components whose transcripts are enriched in cognate Leu codons (Fig. S5A, A’). Overall, our data suggest that fusion protein-mediated increases in TRMT5 are essential for sustaining optimal levels of mitochondrial respiration by enhancing levels of a tRNA target(s), mitochondrial mass, and mitochondrial OXPHOS proteins. Our studies thus provide novel insights into a role for TRMT5 in ARMS tumorigenicity and highlight a mechanism whereby fusion proteins regulate *TRMT5* to enhance mitochondrial OXPHOS activity required to support and sustain ARMS tumors.

### RMS tumor lines exhibit vulnerabilities to inhibition of mitochondrial translation

Our analysis indicated that mitochondrial metabolism is significantly up-regulated in ARMS tumors, and correspondingly, mitochondrially translated OXPHOS proteins were over-produced in a panel of ARMS cell lines and 2D-adapted PDX models (Fig. S6B). Our findings suggest that the underlying mitochondrial translation machinery could prove vulnerable to previously reported inhibitors^51, 56^, such as the antibiotic Tigecycline, which blocks growth of MYC-dependent lymphomas. We therefore tested the impact of Tigecycline, a mitochondrial translation inhibitor, on the viability of two ARMS cell lines, RH4 and CW9019. Importantly, both ARMS cell lines exhibited pronounced sensitivity towards Tigecycline, evidenced by preferential, dose-dependent killing of these cells as compared to normal myoblasts and myotubes (Fig. S6C). The IC_50_ value was found to be ∼2.4 μM for both cell lines based on a normalized response curve (Fig. S6C’). We also tested the impact of Tigecycline on the self-renewal capacity of ARMS cells using colony formation assays. Administration of Tigecycline led to a significant reduction in colony forming units (CFU) in both fusion-positive tumor cells (Fig. S6D-D”), as well as a pronounced reduction in basal and maximal OCR rates in both fusion-positive ARMS cells and 2D-adapted PDX lines (Fig. S6E, E’), indicative of its ability to dampen mitochondrial metabolic activity.

Our *in vitro* data suggest that ARMS tumor cells significantly expand their OXPHOS activity through a concerted, MYCN-independent increase in expression of mitochondrial proteins, rendering their growth susceptible to mitochondrial translation inhibitors.

### RMS tumors display strikingly different therapeutic responses to inhibition of FGFR4 and mitochondrial translation

FGFR4 is a key target of PAX fusion TFs^57^ that is frequently over-expressed in RMS and is associated with aberrant proliferation and metastasis^6, 58^ (Table S5). Interestingly, our chromatin interaction maps uncovered a distinct promoter-anchored interaction with an upstream (∼180 kB) active enhancer, signifying another layer of regulation by the fusion protein resulting in enhanced *FGFR4* expression (Fig. S6F). Our data also indicated that a SE resides immediately downstream of the *FGFR4* oncogene, a known target of PAX3-FOXO1^57^, in cells and O-PDXs expressing PAX7-FOXO1 TF, thereby confirming the reliability of our model in recapitulating early genomic events prevalent in this aggressive form of RMS (Fig. S6F). Further, we reasoned that a therapeutic strategy using a selective, clinically relevant inhibitor of FGFR4, Roblitinib, in tandem with Tigecycline, might synergistically inhibit growth of ARMS tumor cells. We showed that administering Roblitinib significantly inhibited the proliferative and self-renewal capacities of ARMS tumor cells in a dose-dependent manner (Fig. S6G, G’, G”). Moreover, ARMS tumor cells exhibited enhanced sensitivity to combined administration of Roblitinib and Tigecycline as compared to normal myoblasts and myotubes (Fig. S6H) in a synergistic manner *in vitro* (Fig S6I).

We therefore conducted a preclinical phase II study^59^ comprising a panel of O-PDXs to understand the anti-tumor activity of Tigecycline and Roblitinib in animal models of RMS. To recapitulate the heterogeneity of tumors from different RMS patients, we used O-PDXs derived from two PAX3-FOXO1 tumors that were resected at recurrence (SJRHB013759_X14, SJRHB030765_X1), a PAX7-FOXO1 translocated tumor resected at diagnosis (SJRHB013757_X2), and a fusion-negative ERMS tumor resected at recurrence (SJRHB013758_X2) (Table S7). Mice bearing luciferase-labeled O-PDXs were screened weekly using bioluminescence and randomized for enrollment once sufficient luciferase signal (>1×10^6^ photons/sec per cm^2^/sr) was detected or a visible tumor was observed (Table S7). The following seven treatment groups were tested, including single agent arms and combinations using vincristine (VCR) and irinotecan (IRN) as a standard of care backbone: Tigecycline (TIG), Roblitinib (ROB), TIG+ROB, VCR+IRN, TIG+VCR+IRN, ROB+VCR+IRN, and untreated control (Table S7). In total, we enrolled 164 mice and treated them with four courses of therapy (21 days per course) using a clinically relevant schedule (Fig. 6A, Table S7). Mice were monitored weekly for tumor burden, with responses measured by bioluminescence as previously described^59^ (Fig. 6B-G).

**Figure 6.**
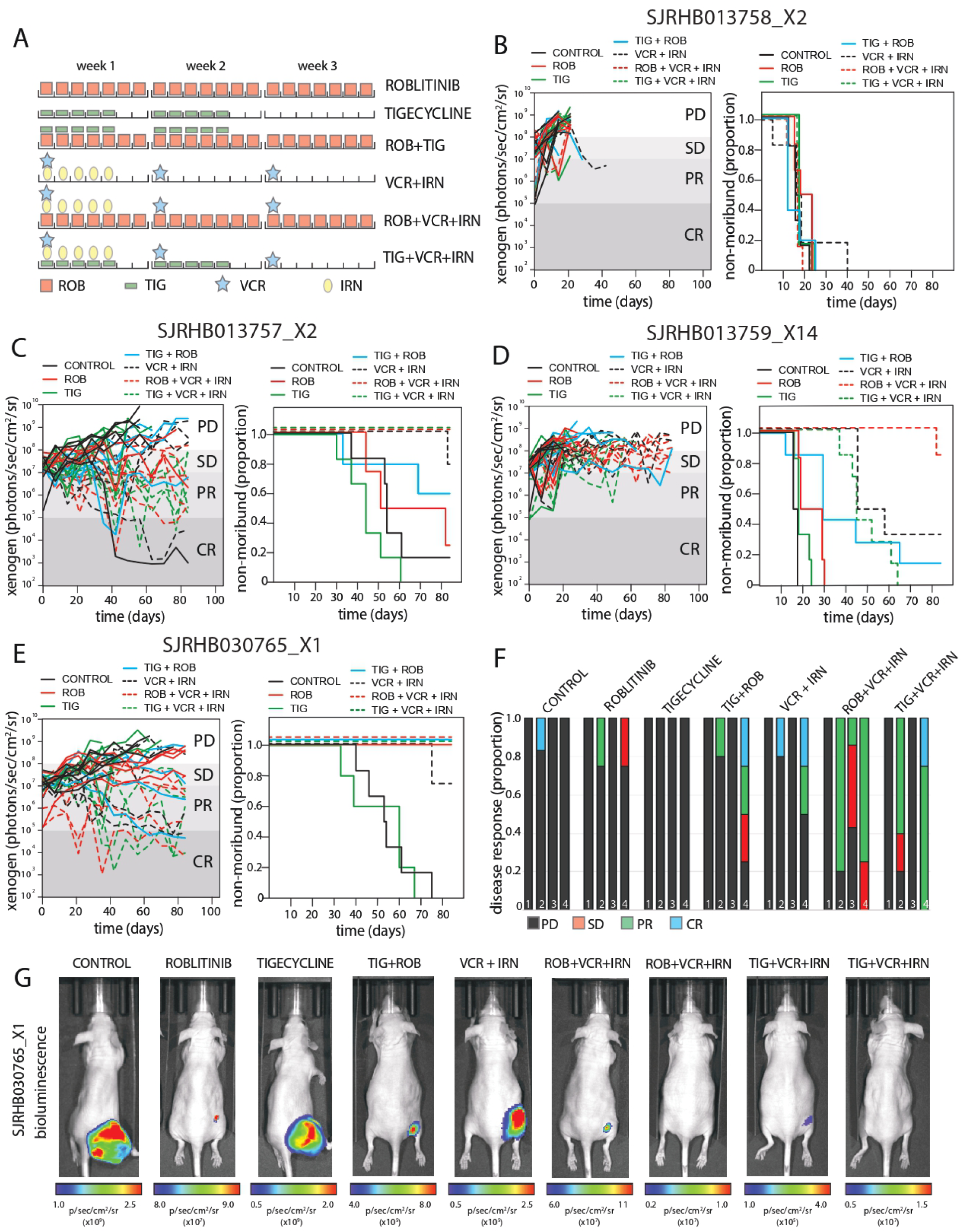
Preclinical phase II testing of roblitinib and tigecycline in RMS O-PDXs. **A)** Schematic of the treatment schedules that were used in preclinical studies for administration of roblitinib (ROB) and tigecycline (TIG) as single agents and in combination with vincristine (VCR) and irinotecan (IRN). **B-E)** Representative survival and individual mouse response for treatment groups shown in **(A)** for the RMS O-PDXS tested: SHRHB013758_X2 (fusion-negative ERMS), SJRHB013757_X2 (PAX7-FOXO1), SJRHB013759_X14 (PAX3-FOXO1) and SJRHB030765_X1 (PAX3-FOXO1). Each O-PDX panel is labeled. **F)** Box plot showing the summary of responses at study endpoint for mice shown in **(B-E).** Bar Numbers: 1, SJRHB013758_X2; 2, SJRHB013757_X2; 3, SJRHB013759_X14; 4, SJRHB030765_X1. **G)** Representative xenogen images of mice from the SJRHB030765_X1 O-PDX preclinical study shown in **(E)**. Abbreviations: PD, progressive disease; SD, stable disease; PR, partial response; CR, complete response. See also Figure S6.

For the fusion-negative ERMS O-PDX (SJRHB013758_X2), no significant responses were seen in single agent ROB or TIG groups or any of the combination arms, and all mice were removed from study with progressive disease by day 40, with an average of 17.9 days on study for all treated groups (Fig. 6B). Remarkably, striking responses were seen in all three of the PAX3/7 fusion-positive O-PDXs (Fig. 6C-F). SJRHB013757_X2 (PAX7-FOXO1) did not have a significant response with ROB or TIG groups as single agents compared to control (p=0.8 and 0.12 respectively). However, TIG + ROB treated mice showed improved survival, with an average of 70.8 days on study compared to TIG alone at 44.5 days (p=0.02; Fig. 6C). SJRHB013759_X14 (PAX3-FOXO1) also had improved survival with TIG+ROB, with an average of 41.9 days on study compared to ROB (avg. 22.5 days) or TIG (avg. 19.5 days) alone (p=0.09 and 0.006 respectively; Fig. 6D). Notably, a strong response in this O-PDX occurred in the triple therapy group of ROB+VCR+IRN with an average survival of 83.7 days on study, which was significant compared to TIG+VCR+IRN (avg. 49.6 days) or VCR+IRN (avg. 60.2 days) treatment groups (p=0.0001 and 0.04 respectively; Fig 6D). The only single agent response was seen in SJRHB030765_X1, with all mice in the ROB group surviving 84 days (p=0.004 and 0.005 compared to untreated control and TIG groups; Fig. 6E). Moreover, mice receiving TIG+ROB, ROB+VCR+IRN, and TIG+VCR+IRN also survived 84 days, with several mice achieving complete responses (by bioluminescence), which were significant compared to TIG alone or untreated control, but not to ROB alone (Fig. 6E-G, Table S7). Overall, mice tolerated treatment well with minimal weight loss or health concerns (Table S7). Taken together, combined treatment with Tigecycline and Roblitinib, a clinically relevant, selective inhibitor of FGFR4, significantly reduced the growth of ARMS tumors in vivo. Interestingly, responses to these drugs varied substantially across the spectrum of RMS sub-types, with significant differences also observed in ARMS tumors harboring each fusion protein.

## Discussion

In this study, we developed a model in which muscle progenitors were programmed to ectopically express PAX fusion proteins that drive development of ARMS. We reasoned that such a system would allow us to study basic mechanisms of oncogenesis in the absence of additional mutations and to identify possible tumor vulnerabilities. The resulting models displayed hallmarks of transformation, and gene expression profiles in these cells revealed key signatures also found in ARMS tumors and O-PDX models, suggesting that these models recapitulated aspects of fusion protein-dependent transformation. It is known that RMS tumors exhibit relatively few recurring cancer-associated mutations^7^, and our data suggest that fusion proteins might bypass the need for additional mutations by activating expression of multiple oncogenes and growth factors. For example, we found that one or both fusion proteins were recruited to super-enhancers that interacted with promoters of *MYCN*, *JUN*, and *FGFR4* (Figs. 2, 3, Table S2). MYCN is frequently amplified in many cancers, including rhabdomyosarcoma^3^ and neuroblastoma, and here, we identified a novel super enhancer that is distinct from previously reported enhancers and the super enhancer found in neuroblastoma^43^, suggesting that assembly of this SE maybe tumor-specific. We found that this fusion-protein-dependent super enhancer interacts with its promoter, in line with enhanced MYCN expression. Activation of this element was also observed in O-PDX samples, suggesting that the MYCN enhancer is of clinical importance for ARMS tumor growth, and indeed, silencing of this region delayed tumor growth in mice. At the same time, oncogenic fusion proteins augment expression of MRFs, *MYOG* and *MYOD1,* which, in turn, up-regulate a large cohort of muscle-specific genes (Fig. S3E, G), but their pro-differentiative abilities are likely counter-balanced by increased expression of inhibitory proteins like Musculin^60^ (MSC) (Table S5) and activities of transcription factors like SIX1^32^, that maintain ARMS tumor cells in an undifferentiated state. Studies with our novel model therefore demonstrate that PAX7-FOXO1 alone is sufficient to up-regulate both MRFs and differentiation-inhibitory factors, endowing RMS cells with characteristics of undifferentiated muscle. Importantly, our iPSC-MP^PAX3/7-FOXO1^ cells will provide a novel resource to explore direct, fusion protein-dependent mechanisms underlying oncogenic transformation.

Our ability to compare isogenic PAX7, PAX7-FOXO1, and PAX3-FOXO1 expressing cells provided additional, novel mechanistic insights into the unique role of PAX7- FOXO1. We found that PAX7-FOXO1 occupied approximately half of the enhancers able to bind PAX7, but the fusion protein had a greater impact on maintaining and promoting gene expression than PAX7, suggesting that replacement of its activation domain with the FOXO1 activation domain significantly boosted expression of muscle signature genes (*MYOD1*, *SIX1*, *SIX4*) and tumor-promoting genes, including MYCN and FGFR4. Further, although we found that both fusion proteins modulate chromatin organization in RMS by promoting long-range promoter-enhancer interactions that drive tumor-specific gene expression, we noted key differences in fusion protein binding events, activation of enhancers and super-enhancers, and expression of target genes in PAX3-FOXO1 versus PAX7-FOXO1 expressing cells (Fig. 2L). Activation of a distinct array of enhancers and super enhancers might contribute to the enhanced aggressiveness of the PAX3-FOXO1 fusion in tumors^6, 9, 10^ (Figs. 2, S2), a possibility that will require additional testing.

Though previous reports have documented the involvement of mitochondrial metabolism in tumor cell migration and invasion^61, 62^, its role in rhabdomyosarcoma growth and proliferation has not been explored. Our data, for the first time, reveal key vulnerabilities of RMS tumor cells with respect to mitochondrial function. We find that RMS tumor cells, on average, have a larger number of mitochondria, and they are distinguished by an enhanced respiratory capacity needed to meet the increased energy demand for tumor growth. Notably, PAX3 fusion-positive tumor cells exhibited higher OXPHOS activity compared to PAX7-fusion cohorts (Fig. 4G). These findings align with a previous report indicating higher mitochondrial OXPHOS activity in Pax3 expressing myogenic cells compared to Pax7 expressing satellite cells^63^. This observation not only reveals key physiological differences between tumor cells expressing each fusion protein but also indicates a heightened mitochondrial response underlying the highly proliferative, undifferentiated state in fusion-positive ARMS tumors. Concordantly, we observed a large number of mitochondrial genes that were up-regulated by both fusion proteins, and this list included numerous proteins from OXPHOS pathway. Our analyses also uncovered *TRMT5* as a key fusion protein target.

The role of TRMT5 has not been thoroughly explored in mammalian cells, and a role in ARMS pathogenesis has not been reported. Our analysis clearly indicates that *TRMT5* plays an essential role in ARMS, as its expression is enhanced in tumors, and it is likely that MYCN collaborates with fusion proteins to drive its expression based on its ability to bind the TRMT5 promoter (mined using dataset^3^). Accumulating evidence^13, 64^ indicates a critical role for tRNAs in driving tumor growth, and our detailed analysis revealed aberrant expression of mitochondrial tRNAs and, remarkably, leucine codon usage bias for genes encoded by the mitochondrial genome. *TRMT5* is a key regulator of the specific iso-acceptor tRNA for leucine (TRNL2), transcripts for which are up-regulated in ARMS tumors. It is likely that other cytosolic and mitochondrial tRNAs, which drive increased mitochondrial protein production, are also aberrantly expressed in ARMS, but they may have escaped detection owing to sub-optimal methods used in RNA-seq studies that fail to capture highly modified tRNAs. Future tRNA-seq and CRISPR-Cas9 gene editing will be needed to explore the role of tRNAs more broadly. Since depletion of TRMT5 led to delayed growth of ARMS tumors, most likely by compromising mitochondrial metabolism, we postulate that therapies based on TRMT5 inhibition could be a novel strategy to treat ARMS tumors. Thus, our data provide a functional link between fusion protein-mediated chromatin reorganization, increased expression of key mitochondrial targets, and reprogramming of mitochondrial metabolism, which drives tumor growth.

Several clinical studies have tested small molecule single-agent and combinatorial treatments for RMS^3, 6, 24, 25^. Unfortunately, due to toxicity, bystander effects, or efficacy issues, most of these drugs were unsuccessful in clinical or preclinical stages, prompting the need to discover novel therapies. Further, FGFR4-based monotherapy frequently leads to therapy resistance in many cancers, including rhabdomyosarcoma^65, 66^, prompting the need for combinatorial therapies. Our findings indicating over- expression of mitochondrial proteins and FGFR4 in ARMS led us to examine vulnerabilities of ARMS tumors to small molecule inhibitors of these targets, and we formulated a novel strategy that combined Tigecycline with a clinically relevant FGFR4 inhibitor, Roblitinib, whose clinical efficacy in treating ARMS has not been explored. Roblitinib displays excellent specificity toward FGFR4 by interacting with a cysteine residue in the FGFR4 active site that is not present in other FGFRs^67^. Moreover, since both compounds display outstanding pharmacokinetic and safety properties in mouse studies and clinical trials for hepatocellular carcinoma (Roblitinib)^68, 69^ and leukemia (Tigecycline)^56, 70^, we tested these compounds in mice. Remarkably, we found that combined administration of Tigecycline and Roblitinib significantly inhibited ARMS tumor cell growth, viability, and mitochondrial metabolism using *in vitro* and *in vivo* O-PDX models. Most importantly, our pre-clinical phase II data evidently confirms the potential of the drug combination therapy with TIG and ROB module ROB+TIG to offer significant survival benefits and complete responses (CR) in mice bearing PAX3-fusion tumors, with several mice showing durable responses of at least 8 weeks following the completion of therapy. Interestingly, we also observe differential drug responses for PAX3 and PAX7-fusion tumors, a finding that requires incorporation of additional models and testing. Nevertheless, ROB+VCR+IRN and TIG+VCR+IRN were found to be highly efficacious against both PAX-fusion tumors, thereby signifying the utility of these treatments for future clinical trials in patients with high-risk RMS. Collectively, our preclinical phase II study represents an outstanding starting point to validate additional therapeutic combinations in a larger cohort of preclinical models in future trials. Moreover, our novel therapeutic approach targeting FGFR4 and mitochondrial functions might be broadly applicable to patients with solid tumors of liver, cholangiocarcinoma, kidneys, and colon that display aberrant *FGFR4* activity^71–74^.

## Acknowledgements

We thank Arima Genomics for providing an award to BDD that included their pCHiC kit. This work was supported by a grant from the NIH (R01 AR 076954) to BDD. BK is funded by a grant from The New York State Stem Cell Science Program (NYSTEM, #C322560GG). ES and MAD are supported by the National Cancer Institute (U01CA263969) and ALSAC. Work by ES in this publication is supported by the Oncology Center of Excellence, Food and Drug Administration (FDA) of the U.S. Department of Health and Human Services (HHS) contract number 75F40121C00092. This work was supported by an NCI Cancer Center Support Grant (CA21765). MAD received funding support through the NIH grants CA219686, CA248432, and CA24550. We thank Brittney Gordon, Kaley Blankenship and Asa Karlstorm, Departments of Oncology and Developmental Neurobiology, St Jude Children’s Research Hospital for assisting with the pre-clinical studies. O-PDX models were provided from the Childhood Solid Tumor Network, also from St. Jude. We thank Dr. J. Khan (NIH) for providing PFM2.1 antibody. We thank Dr. M. Thangavel, Neuroscience Institute, NYU Grossman School of Medicine, for assisting with animal studies at NYU animal facility. We thank Dr. T. Rodrick at the Metabolomics Core Resource Laboratory, for assisting with Seahorse assays, and the NYU School of Medicine Genome Technology Core. We also thank Ilya Shamovsky, Department of Biochemistry and Molecular Pharmacology, NYU Grossman School of Medicine, for assisting with the NGS experiments. We thank A. Liang, J. Sall, and C. Petzold at the Imaging Core for assistance with TEM. We are grateful to Dr. A. Cherkasov (U. British Columbia) for providing us with VPC-70619. The authors thank the Children’s Oncology Group Soft Tissue Sarcoma Committee and the BioPathology Center, for their careful collection of clinical samples. This research was supported by the Intramural Research Program of the National Institute of Health and National Cancer Institute. BDD is most grateful to Davide Ruggero and Kevan Shokat (UCSF) for inspiring discussions.

## Author contributions

Conceptualization, B.K., E.S., and B.D.D.; Methodology, B.K., G.M.C., J.M., M.A., M.A.D., and E.S.; Investigation, B.K., G.M.C., J.M., M.A., M.K., A.M. and E.S.; Formal Analysis, G.M.C, B.K. and E.S.; Writing – Original Draft, B.K.; Writing – Review & Editing, B.K., G.M.C., J.M., E.S., and B.D.D.; Funding Acquisition, B.K., E.S., M.A.D., R.C.R.P. and B.D.D; Resources, M.A.D., E.S., R.C.R.P. and B.D.D; Supervision, M.A.D., E.S. and B.D.D.

## Supplementary Information

Document S1: Figures S1-S6.

Table S1: Excel file containing analyzed RNA-seq data from iPSC-MP ^PAX3/7FOXO1^ cells.

Table S2: Excel file containing analyzed pCHi-C data from iPSC-MP ^PAX3/7FOXO1^ cells.

Table S3: Excel file containing analyzed Hi-C/TADCompare data from iPSC-MP PAX3/7FOXO1 _cells._

Table S4: Excel file containing GO terms for mitochondrial transcriptional targets of PAX3-FOXO1 and PAX7-FOXO1 fusion TFs.

Table S5: Excel file containing differential gene expression value in iPSC-MP ^PAX3/7FOXO1^ cells, primary ARMS tumors and O-PDX samples.

Table S6: Excel file containing genome-wide binding data for PAX3-FOXO1 and PAX7- FOXO1.

Table S7: Excel file containing preclinical testing data for Tigecycline, Roblitinib, Vincristine and Irinotecan in RMS O-PDX.

## Methods

### Cell Culture

Human iPSC cells expressing PAX7 (muscle progenitors)^21, 22^ were grown on 0.1% gelatin-coated culture plates in GlutaMAX supplemented IMDM (Gibco) with 15% stem cell qualified fetal bovine serum (Gemini), 1% penicillin/streptomycin (Corning), 50 μg/ml L-ascorbic acid (Sigma), 4.5 mM 1-thioglycerol (Sigma), and 10 ng/ml recombinant human basic-FGF (Peprotech). These muscle progenitors were infected using lentiviral doxycycline-inducible (pTRE-PURO) or constitutive (EF1α promoter-PURO) constructs driving expression of a cDNA encoding the exact PAX7-FOXO1 or PAX3-FOXO1 fusion protein found in ARMS tumors. PAX7 transgene expression was induced by adding Dox (Sigma-Aldrich, final concentration 1 μg/ml), and then maintained throughout the protocol by replacing the medium (including dox) with 10 ng/ml human basic-FGF (Peprotech) every 2 days. Cells were expanded using the same media with Dox for 3 or 4 additional days before harvesting them for analysis. For cultures without Dox, cells were harvested at day 3 following Dox removal for analysis.

Human alveolar rhabdomyosarcoma (ARMS) cell lines SJRH30 (RH30) and RH4 were gifts from Prof. Pier-Luigi Lollini (University of Bologna, Italy). CW9019 was a gift from Dr. J. Khan (NIH, Bethesda, US). RH30 and CW9019 cells were cultured and maintained in Dulbecco’s Modified Eagle Medium (DMEM) (Thermo-Fisher Scientific, USA; Cat# 11995065) supplemented with 10% (v/v) fetal bovine serum (FBS) (Sigma, USA; Cat# F2442) and 1% penicillin-streptomycin (Gibco; Cat#15140122) at 37°C, 5% CO2 and 95% humidity. RH4 cells were cultured and maintained in RPMI 1640 Medium (Thermo-Fisher Scientific, USA; Cat# 11875093) supplemented with 10% (v/v) fetal bovine serum (FBS) (Sigma, USA; Cat# F2442) and 1% penicillin-streptomycin (Gibco; Cat#15140122) at 37°C, 5% CO2 and 95% humidity. Human HEK cells were cultured and maintained in DMEM supplemented with 10% (v/v) FBS and 1% penicillin-streptomycin at 37°C, 5% CO2 and 95% humidity. Human primary skeletal muscle myoblasts (Lonza, HSKMM, Cat# CC-2580) were cultured according to the supplier’s protocols.

### RNA-seq

Total RNA was extracted from iPSC control and iPSC-MP^PAX3/7-FOXO1^ cells (in three replicates) using RNA Zymo-Spin columns (Zymo R1015). Before library preparation, rRNA was depleted with NEBNext rRNA Depletion Kit (NEB E6310). RNA-seq libraries were constructed using the NEBNext Ultra II Directional RNA Library Prep Kit for Illumina (NEB E7760) according to the manufacturer’s instructions. Libraries were sequenced (2x50bp paired end) using an Illumina NextSeq 2000 machine. Raw sequencing reads were aligned to the human reference genome (hg38 assembly) with bowtie2 v2.3.5.1^75^. Quantification of gene expression was performed with featureCounts from Subread v1.6.3 using uniquely mapping reads^76^. Subsequent analysis was done in R using the DESeq2 package to calculate differentially expressed genes^77^. To account for significant differences, we considered genes to be up-regulated or down-regulated if the fold- change in expression in the model cell lines was ≥ 2 with an FDR value < 0.05. We considered the remainder of the genes unchanged between cell lines. For GSEA, we used the R package clusterProfiler, selecting for ontologies related to biological processes (BP) with a minimum of 10 genes and a p-value cutoff of 0.05^78^. In order to standardize our analysis and to compare it with our own data, we analyzed RNA-seq counts from 26 primary ARMS tumors^7^ and normal myoblasts^79–81^ as described for our study. The lists of differentially expressed genes were compared to obtain overlaps between our cell lines and tumors. tRNA expression and quantification was done using the R package coRdon for the genes expressed for each cellular model^82^.

### Cut&Run Seq

Cut&Run was performed on 0.25 x 10^6^ cells, with two replicates per condition, using the Cell Signaling Cut&Run kit (#86652) according to the manufacturer’s instructions. We used 2 μg of antibody or normal rabbit IgG as background control with overnight incubation at 4°C (a list of the antibodies used can be found in Key Resource Table). Protein A/G coupled MNase was incubated with antibodies to digest proximal DNA. Fragmented DNA was extracted and purified using DNA spin columns (Cell signaling #14209). The enriched DNA was used to construct sequencing libraries with NEBNext Ultra II DNA Library Prep Kit for Illumina (NEB E7103). Libraries were sequenced (2 x 50bp paired end) in an Illumina NextSeq 2000 machine. Raw reads were aligned with bowtie2 v2.3.5.1 to the human genome (hg38 assembly)^75^. Next, we removed unmapped and duplicate reads with MarkDuplicates from Picard v2.18.11^83^ and called peaks using MACS2 v2.1.1 (the --broad option was used for H3K4me1)^84^. BigWig files were obtained using deeptools v3.2.1^85^ and bedtools v2.26.0^86^.

Enhancers were obtained by overlapping H3K27ac and H3K4me1 peaks and excluding promoter regions with H3K4me3 signal. Super enhancers were computed with ROSE software^30^ using H3K27ac BAM files and the previously computed enhancer list as input. Peak overlaps between cell lines were obtained using bedtools intersect^86^. De novo motif analysis was performed using HOMER^87^ for the central 500 bp of the top 5000 peaks per condition and then compared to known motifs. Peak annotation was also done using HOMER software.

### HiC and promoter capture HiC (pCHiC)

For HiC and pCHiC, we used a commercial kit from Arima Genomics (A510008). Briefly, 10^6^ cells were cross-linked with formaldehyde and extracted following the manufacturer’s instructions. 4 μg of DNA per condition were digested and ligated, and biotin was incorporated to select successfully ligated fragments. Ligated DNA was sonicated with a Diagenode Bioruptor for 20 min, 30 seconds ON/OFF, at 4°C to obtain ∼400 bp DNA fragments. Biotinylated DNA was purified using streptavidin beads. Next, 200 μg of sonicated and ligated DNA was used as input for library preparation using ARIMA’s indexed primers. A fraction of the HiC library was used for sequencing in an Illumina NovaSeq 6000 for 300 cycles (paired-end) at a depth of ∼200M reads per sample.

Next, the remainder of the HiC library was used for promoter capture using ARIMA’s custom panel of human promoters (∼84,000 probes for ∼26,000 promoters). After promoter capture, the pCHiC libraries were amplified following the manufacturer’s instructions. The promoter capture libraries were sequenced with an Illumina NovaSeq 6000 (300 bp, paired-end) at a depth of ∼200M reads per sample.

### Data processing for HiC

Raw sequencing reads were processed using juicer v1.4^88^ with genome assembly hg38 and ARIMA’s restriction sites. Replicates for each cell line were merged after alignment within the juicer pipeline. The resulting HiC files containing contact matrices were used to calculate the first principal component (PC) with juicertools v2.20.00 Eigenvector at 250k resolution with VC normalization. The A and B compartments were established by correlating the PC sign with H3K27ac for each cell line. The hic files were also used to compute TADs with juicertools v2.20.00 arrowhead with VC normalization at 25k resolution. R packages strawr^88^ and TADCompare^89^ were used to compute differences in TAD boundaries between cell lines.

### Data processing for pCHiC

Raw sequencing reads were processed using ARIMA’s pipeline for promoter capture experiments (Arima-CHiC-v1.4). Briefly, we used the HiCUP^90^ pipeline from the Babraham Institute to map with bowtie2 each pair of the ligated fragments to the human genome (hg38 assembly). Next, BAM files were converted into CHiCAGO input files and processed with CHiCAGO^91^ to obtain interaction confidence scores for each cell line. Only cis- and significant interactions (interaction scores ≥5) were considered. These interactions were mapped to a custom list of enhancers for each cell line (from our Cut&Run analysis) to obtain a list of significant promoter-enhancer interactions. We used the WashU epigenome browser to simultaneously visualize pCHiC interactions and other sequencing data^92^.

### TCGA data analysis

TPM normalized counts from the TCGA datasets were obtained from the Genomic Data Commons portal (https://portal.gdc.cancer.gov/) and compared to TPM normalized counts for ARMS tumors. Chronos dependency scores from CRISPR screening (DepMap Public 23Q4+Score, Chronos) for the indicated genes were obtained from the DepMap portal https://depmap.org/portal/.

### RNA isolation and quantitative PCR (qPCR)

Total RNA was extracted from cells using Invitrogen™ TRIzol™ Reagent (Fischer Scientific, Cat# 15-596-018) followed by phenol-chloroform purification. cDNA synthesis was performed using Verso cDNA synthesis kit (Thermo Fisher, Cat# AB1453A). The cDNA samples were then used as templates for qPCR on an ABI 7500 Fast Real-Time System (Applied Biosystems) using Power SYBR™ Green PCR Master Mix (Thermo Fisher, Cat# 4367659) and specific primers for each gene (Key Resource Table).

### Immunoblotting

Protein lysates were prepared from cells that were plated, cultured, and harvested at indicated time points as described above. Briefly, cells were lysed in ice-cold ELB lysis buffer (1M Hepes pH 7, 5M NaCl, 500 mM EDTA, 0.1% NP-40, 10% glycerol, and 1 mM DTT) containing a protease/phosphatase inhibitor cocktail (AEBSF, leupeptin, aprotinin, NaF, β-glycerophosphate). Equal amounts of protein were separated by SDS-PAGE and transferred to PVDF membrane (Millipore; Billerica, MA, USA; Cat# iPVH00010). Western blotting was performed using standard procedures, with blocking in 5% milk (Himedia; Cat# RM1254-500GM) for 3 to 5 hours, washing in PBS containing 0.2% Tween 20 (Sigma; Cat# P7949-100ML), incubation overnight in primary antibody at 40C, and 2hour incubation with secondary antibody at room temperature. HRP-conjugated secondary antibodies were used, signal was detected using the Thermo Scientific™ SuperSignal™ West Dura Extended Duration Substrate (ThermoFisher; Cat# PI34076), and blots were exposed to film. Antibodies and relevant dilutions are listed in Key Resource Table.

### Cell survival assay

The effect of Tigecycline, Roblitinib, or Tigecycline+Roblitinib on the viability of RH4, CW9019, HSMM (myoblast), and human myotubes was examined using the MTT assay. Briefly, 10,000 cells were plated in flat-bottomed 96-well microplates and cultured overnight as described above. The cells were incubated the following day with different concentrations of Tigecycline, Roblitinib, or Tigecycline+Roblitinib for 48 hours. Thereafter, the medium was replaced with 3-(4,5-dimethylthiazol-2-yl)-2,5- diphenyltetrazolium bromide (MTT, Sigma-Aldrich) dissolved at a final concentration of 1 mg/ml in serum-free, phenol red-free medium. Following incubation of 2 hours, the formazan crystals were then dissolved by DMSO and the absorbance of the solution was measured at 570 nm. Survival of DMSO-treated control cells was set at 100%.

### Clonogenic Assays

For all anchorage-dependent colony formation assays, cells were plated in 6-well dishes at 4000 cells/well. For drug studies, RH4 and CW9019 cells were incubated the following day with different concentrations of Tigecycline and Roblitinib for 48 hours and cultured for 9 days. Colonies were washed twice with ice cold 1×PBS, fixed in 10% neutral buffered formalin (30 min), and subsequently stained with 0.5% crystal violet (∼30 min). Respective wells were imaged using GelDoc, and colony forming units or colonies comprising >50 cells were counted manually under a stereomicroscope. Clonogenic assays were performed independently a minimum of three times, and graphical data are represented as mean ± SEM.

### Tumor Sphere Assay

*In vitro* tumor sphere assays were performed using published methods^93^ with minor modifications. iPSC-MP controls and myogenic progenitors expressing fusion proteins (iPSC-MP^PAX3/7-FOXO1^) cells were grown in defined media +/-Dox until 80-90% confluency was reached. After decanting the media, cells were scraped with a sterile cell scraper and collected in media (DMEM/F12, 1 % Penn-strep, 1% N2 supplement, 10ng/ml hEGF, 10ng/ml β-FGF). Thereafter, cells were seeded at a density of ∼200 cells/well in Nunc low attachment 96-well plates. Cells were then scored for tumor sphere formation under a phase contrast microscope at day 7.

### CRISPRi Interference (CRISPRi)

For silencing regulatory regions (promoters and enhancers) with CRISPRi, we designed 20 bp sgRNAs against the annotated regions (based on H3K4me3, H3K27ac, PAX3/7- FOXO1, or H3K4me1 CUT&RUN signal) using a publicly available portal CRISPick (http:// https://portals.broadinstitute.org/gppx/crispick/public). The detailed methodology for cloning sgRNA oligonucleotides into a lentiGuide-Zeo vector was previously described^31^. RH4 and CW9019 cells were infected with lentiviruses expressing dCas9- KRAB-MeCP2 (a kind gift from N. Sanjana, New York Genome Center), and selected with blasticidin (10 μg/ml) for a week. Medium was replaced with selection media every 48 h until all negative control cells were dead. Following blasticidin selection, cells were transduced with different sgRNAs and selected with Zeocin (1 mg/ml), changing media every 48 h until all negative control cells were dead. The list of sgRNA oligonucleotides we employed can be found in Key Resource table.

### tRNA expression

Total RNA was extracted from cells using Invitrogen™ TRIzol™ Reagent (Fischer Scientific, Cat# 15-596-018) followed by chloroform extraction and elution in RNase/DNase free H_2_O. ∼5μg of the extracted O-PDX RNA was processed for tRNA pretreatment and first-strand cDNA synthesis using rtStar™ tRNA Pretreatment & First-Strand cDNA Synthesis Kit (Arraystar Inc., Cat# AS-FS-004) according to manufacturer’s protocol. Individual tRNA expression was normalized to U6 and was scored using rtStar™ Pre-designed Human tRNA Primer Set (Arraystar Inc., Cat#AS- NR-001H-1-175).

### Flow Cytometry

Mitochondrial membrane potential and mitochondrial mass were measured in ARMS cell lines (RH4, CW9019) and human myoblasts using TMRM (Tetramethylrhodamine methyl ester perchlorate) and Mitotracker Red uptake. Briefly, 0.1x10^6^ cells were incubated with TMRM (250 nM) or Mitotracker Red (50 nM) (# M7512) at 37°C for 30 mins or 2 hrs, respectively. Cells were washed twice with PBS and resuspended in FACS analysis buffer. Data were acquired using a FACS Fortessa instrument (Becton Dickinson Biosciences, USA).

### Generation of expandable 3D and 2D-adapted PDX models

2D-adapated cells were derived from 3D spheroid models cultured from rhabdomyosarcoma orthotopic xenograft (O-PDX) tumor cells. To generate 3D tumor spheroids, O-PDX tumor cells were seeded onto 96-well low-adhesion U-bottom plates (Thermo, MS-9096UZ) at a density of 50,000 cells/well. Plates were centrifuged at 300*xg* for 3 minutes to form centralized cell pellets in each well. Spheroids form after 5- 7 days and were then transferred to 5% geltrex-coated (Thermo, A14133-02) 10 cm tissue culture plates. Within 1-2 days, spheroids attach to the dish and cells migrate out of the spheroid and proliferate. After 5-7 days, cells were passaged one time (passage 1) to expand for cryopreservation. 2D-adapted lines were then thawed on 5% geltrex-coated dishes with appropriate medium supplemented with 1:1000 Thiazovivin (Stem Cell #72254) and tested for viable passaging, and mouse cell contamination. Short tandem repeat (STR) analysis was used to validate original patient STR profile. 3D and matched 2D cells were cultured in either basal medium (BF) or basal medium supplemented with 10% fetal bovine serum. BF medium contains SkGM-2 (Lonza cat # 3246 medium kit) supplemented with L-glutamine, rhEGF, dexamethasone, GA-1000, 1X B-27 (Gibco cat. # 12587010), 1:1000 heat stable bFGF (Gibco, Cat. # PHG0369), and 1:1000 Heparin (Stem cell technologies Cat. # 07980). For GF medium, 10% FBS was added to the BF medium.

| 2D model | Subtype | Fusion Status | Medium |
| --- | --- | --- | --- |
| SJRHB013757_C1 | ARMS | PAX7/FOXO1 | BF |
| SJRHB013759_C2 | ARMS | PAX3/FOXO1 | BF |
| SJRHB013759_C3 | ARMS | PAX3/FOXO1 | GF |
| SJRHB031320_C1 | ARMS | PAX7/FOXO1 | GF |
| SJRHB030765_C1 | ARMS | PAX3/FOXO1 | BF |

### Seahorse Assays

Mitochondrial respiration was measured in representative ARMS cells (RH4, CW9019), 2D-adapted PDX models (SJRHB013757_C1, SJRHB013759_C2),human myoblasts (HSMM), iPSC-MP, iPSC-MP^PAX3FOXO1^, iPSC-MP^PAX7FOXO1^, MYCN depleted CW9019 and RH4 cells using Agilent Seahorse XF Cell Mito Stress Test assay kit (#103015-100) following the manufacturer’s recommendations. Briefly, cells at 70-90% confluency were plated in 24-well Seahorse plates. For Tigecycline experiments, cells were pre-treated with Tigecycline for 6 hr prior to assay. Cells were washed twice with Seahorse assay media before incubating in an incubator without CO_2_ for 1 hr. Following incubation, cells were treated in succession with Oligomycin (1.5μM), FCCP (0.5μM), and a Rotenone/Antimycin A (0.5μM) mixture, and oxygen consumption rate (OCR) was measured in the medium using a Seahorse Analyzer XF/XF24 analyzer.

### Transmission Electron Microscopy (TEM)

Briefly, cells were fixed with a 0.1 M phosphate buffer (pH 7.3) containing EM-grade 2% paraformaldehyde, 2.5% glutaraldehyde, 2mM CaCl2 in 0.1M sodium cacodylate buffer at room temperature. Cell pellet collected were washed with 0.1M sodium cacodylate buffer pH 7.2 at 4°C. Following, 1% osmium tetroxide (OsO_4_) + 1% potassium ferrocyanide (K_4_Fe(CN)_6_) in 0.1M sodium cacodylate buffer were added and incubated for 1 hr at 4°C, and washed with 0.1M sodium cacodylate buffer at 4°C. Thereafter, cells were washed twice with ddH_2_O at 4°C and incubated overnight in uranyl acetate 0.25% at 4°C. Next, dehydration was performed in a graded ethanol series, followed by infiltration with propylene oxide and epoxy resin sequentially, at room temperature. After transferring sample pieces into embedding molds, resin polymerization was performed at 60°C for 2 days. Sections were collected and stained with 3% uranyl acetate in 50% methanol and lead citrate. Images were captured on a JEOL1400 Flash transmission electron microscope.

### Incucyte Label-free Cell Confluency Assay

Briefly, RH4 cells were seeded at 3000 cells/well onto a 96-well tissue culture plates. For drug studies, cells were incubated the following day with different concentrations of Tigecycline (5μM, 10μM) and Roblitinib (1μM) individually, as well as in combination (ROB+TIG) treatment regimen. Thereafter, the cell plates were placed into the Incucyte Live-Cell Analysis System. Pre-warming to 37°C for 30 minutes was performed prior to scanning. Scans were performed for a total of 96 hrs and analyzed using the Incucyte Cell-by-Cell Analysis Software Module.

### Animal Experiments

#### Animals

Athymic nude immunodeficient mice were purchased from Charles River Laboratories (strain code 553). This study was carried out in strict accordance with the recommendations in the Guide to Care and Use of Laboratory Animals of the National Institute of Health. The protocol was approved by the Institutional Animal Care and Use Committee at St. Jude Children’s Research Hospital. All efforts were made to minimize suffering. All mice were housed in accordance with approved IACUC protocols. Animals were housed on a 12-12 light cycle (light on 6 a.m./off at 6 p.m.) and provided food and water *ad libitum*.

For human RMS cell line xenograft experiments, 5x10^6^ cells were resuspended in PBS, mixed 1:1 with Matrigel (Corning, Cat# 356234) and injected subcutaneously between the flank and mid-abdomen region of 6 to 8-week-old female NOD/SCID or NSG mice. Tumor volumes were determined using the formula width^2^ × length × 0.5. Measurements were made weekly until the terminal end-points of the study were reached.

#### Phase II Preclinical Trials

Rhabdomyosarcoma orthotopic xenografts were created by injecting luciferase labeled tumor cells, dissociated and passaged according to the MAST protocol (NCT01050296), into recipient athymic nude mice using an intramuscular injection technique. 4 xenograft lines with the following demographics were tested:

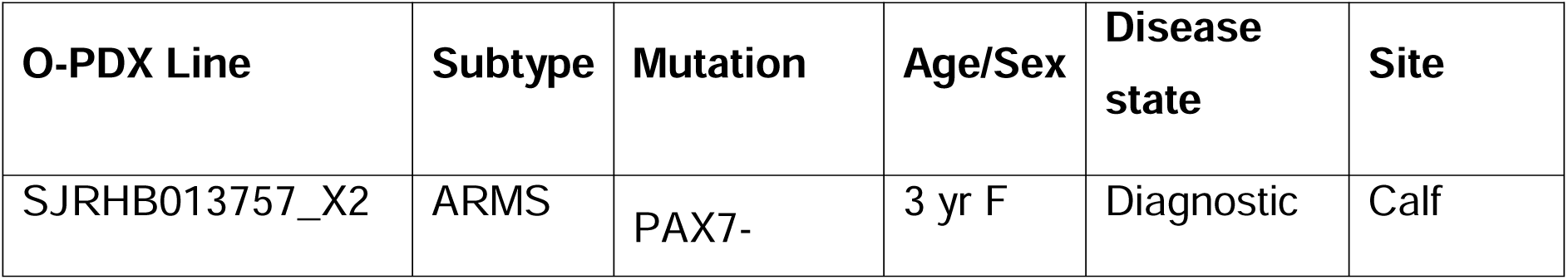
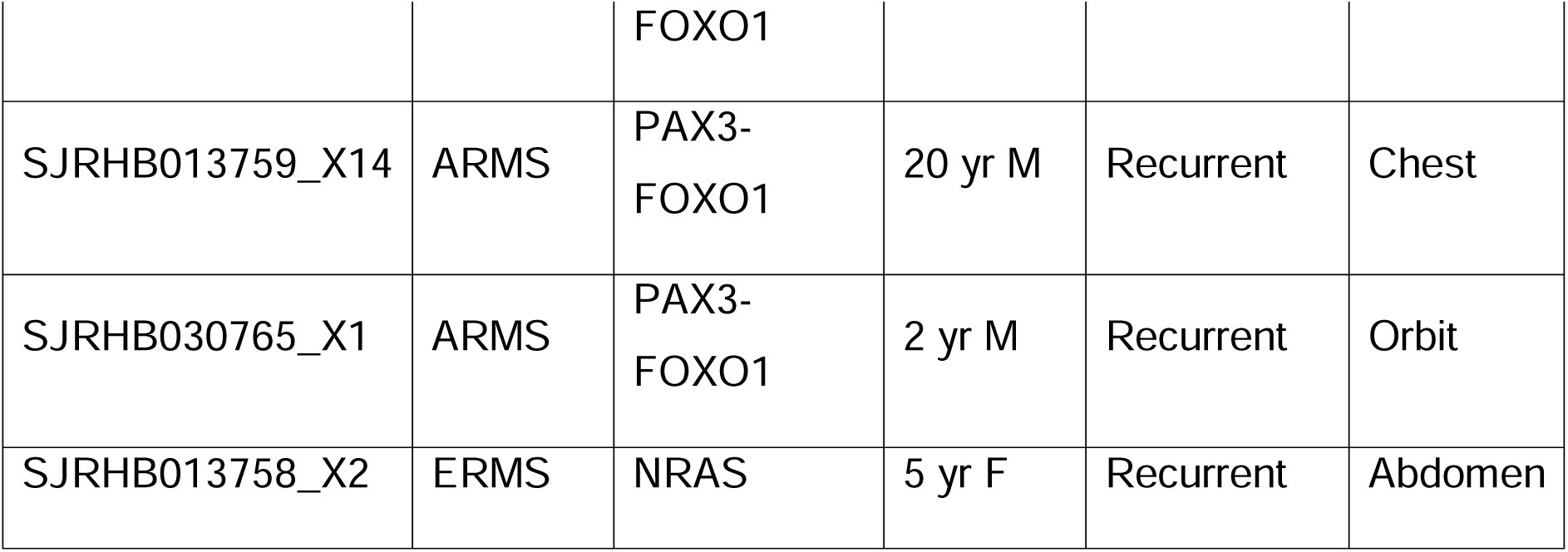

Mice were screened weekly by Xenogen imaging, and bioluminescence was measured. Mice were enrolled in the study after achieving a target bioluminescence signal of 10^7^ photons/sec/cm^2^ or a palpable tumor, and chemotherapy was started the following Monday.

Each xenograft line was separately randomized to the following 7 treatment groups: Control, Tigecycline, Roblitinib, Tigecycline + Roblitinib, Vincristine + Irinotecan, Tigecycline + Vincristine + Irinotecan, Roblitinib + Vincristine + Irinotecan

The following doses and schedules were used for each drug:

|  |  |
| --- | --- |
| Tigecycline | 50 mg/kg oral gavage once daily on days 1-5 and 8-12 |
| Roblitinib | 30 mg/kg oral gavage twice daily on days 1-21 |
| Vincristine | 0.25 mg/kg intraperitoneal once daily on days 1, 8, and 15 |
| Irinotecan | 1.6 mg/kg intraperitoneal once daily on days 1-5 |

Mice received 4 courses of chemotherapy (3 weeks per course), and bioluminescence was monitored weekly and at the end of therapy. Disease response was classified according to bioluminescence signal. Mice with a signal of less than 10^5^ photons/sec/cm^2^ (similar to background) were classified as complete response, 10^5^-10^6^ photons/sec/cm^2^ as partial response, 10^7^-10^8^ photons/sec/cm^2^ (similar to enrollment signal) as stable disease, and greater than 10^8^ photons/sec/cm^2^ as progressive disease. Mice with tumor burden at any time greater than 20% of body weight were removed from the study and classified as progressive disease. Mice were monitored daily while receiving chemotherapy.

### Xenogen Imaging and Quantification

Mice injected with the luciferase labeled cell lines were given intraperitoneal injections of Firefly D-Luciferin (Caliper Life Sciences 3 mg/mouse). Bioluminescent images were taken five minutes later using the IVIS® 200 imaging system. Anesthesia was administered throughout image acquisition (isoflurane 1.5% in O_2_ delivered at 2 liters/min). Living Image 4.3 software (Caliper Life Sciences) was used to generate a standard region of interest (ROI) encompassing the largest tumor at maximal bioluminescence signal. The identical ROI was used to determine the average radiance (photons/s/cm^2^/sr) for all xenografts.

### Data availability

All the raw data and processed files from the RNA-seq, Cut&Run, HiC and pcHiC generated within this manuscript are deposited under the series reference GSE253895 (Token: ixqbaoiqrjgnjkh). Human Pax7 binding data from ChIP-seq was obtained from GSE98976. PAX3-FOXO1 ChIP-seq binding data in RH4 cell lines was obtained from GSM2214084 and GSE163068. O-PDX RNA-seq and ChIP-seq data for ARMS and ERMS tumors was obtained from St Jude’s Children Hospital Cloud within the Childhood Solid Tumor Network (CSTN)^94^. ARMS tumor RNA-seq and ChIP-seq data from patients was obtained from the database of Genotypes and Phenotypes (dbGaP) study accession: phs000720.v4.p1. Human normal myoblast RNA-seq data was obtained was obtained from GSE117609, GSE78649 and^79^.

## Supplementary Information

**Figure S1.**
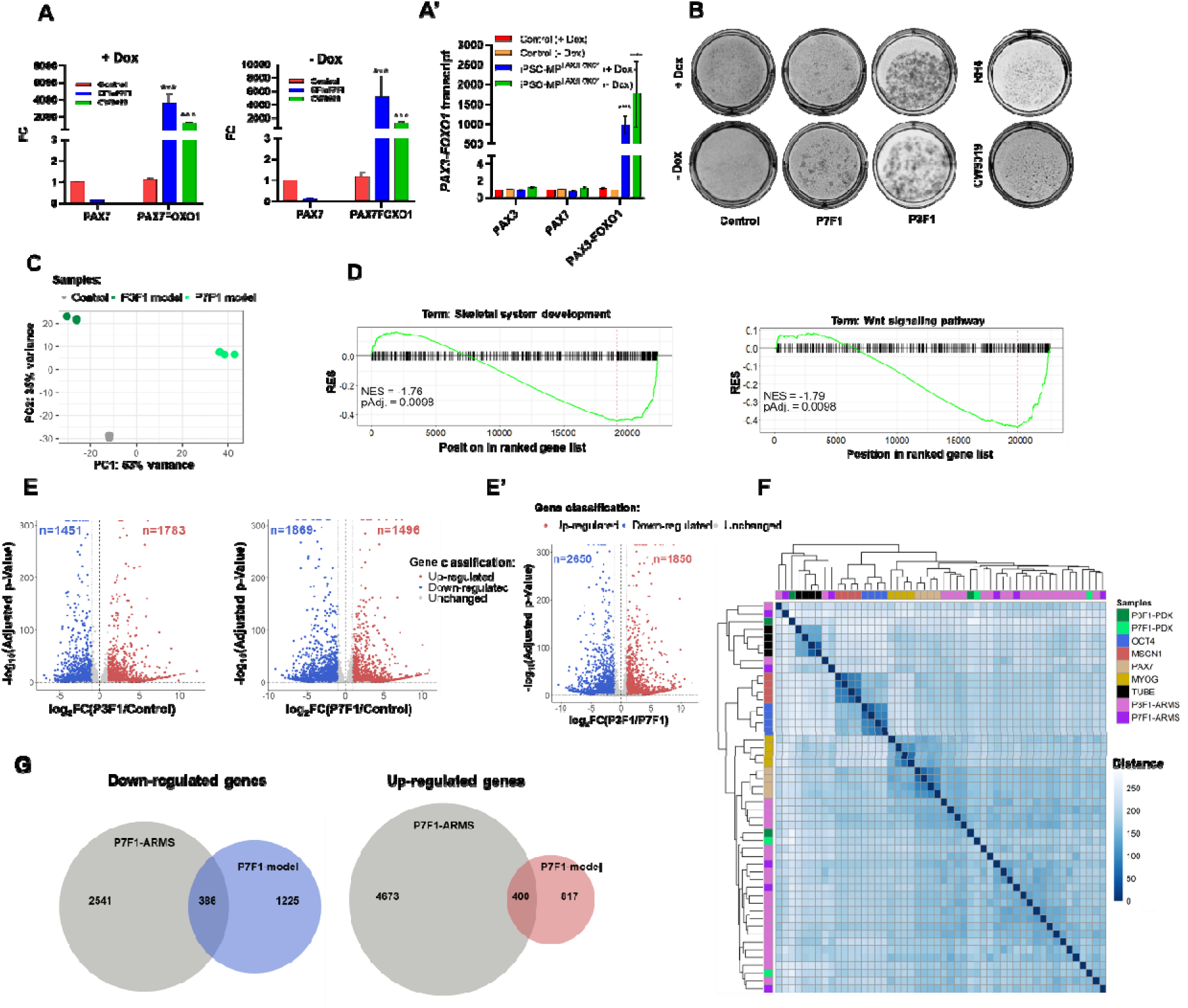
Investigating growth characteristics of oncogenic human PAX-fusion myogenic progenitor model. **A)** *PAX7-FOXO1* transcripts in human myogenic progenitors expressing PAX7-FOXO1 (EF1α P7F1), fusion-negative control cells, and CW9019 +/- Dox. **A’)** *PAX3-FOXO1* transcripts in human myogenic progenitors expressing PAX3-FOXO1 (iPSC-MP^PAX3-FOXO1^) and fusion-negative control cells at +/- Dox. n = 3 per group; error bar, SEM; ***p < 0.001, ****p < 0.0001, ANOVA, Dunnett’s test. **B)** Representative images depicting colony forming micrographs comparing clonogenic growth potential of iPSC-MP^PAX3/7-FOXO1^ with their respective representative cell lines, RH4 and CW9019 respectively at +/– doxycycline condition. **C)** Principal component analysis (PCA) of transcriptomic data in cells expressing PAX3-FOXO1 (left and PAX7-FOXO1 (right) compared to fusion negative controls, following removal of Doxycycline (-Dox). **D)** Gene-set enrichment analysis (GSEA) of transcriptomic data from iPSC-MP ^PAX7FOXO1^ showing Running Enrichment Score (RES) of selected gene ontology terms. **E)** Volcano plots for differentially expressed genes in iPSC-MP cells expressing PAX3-FOXO1 and PAX7-FOXO1 transcription factors. **E’)** log_2_FC plot for genes down/up-regulated in iPSC-MP^PAX3-FOXO1^ versus iPSC-MP^PAX7-FOXO1^ cells. **F)** Sample distance mapping matrix comparing gene expression profile of different muscle lineage cells with alveolar rhabdomyosarcoma (ARMS) tumors and O-PDX. **G)** Venn diagram depicting overlapping changes in transcriptomic data for PAX7-FOXO1 positive ARMS tumors and iPSC-MP^PAX7-FOXO1^ cells.

**Figure S2.**
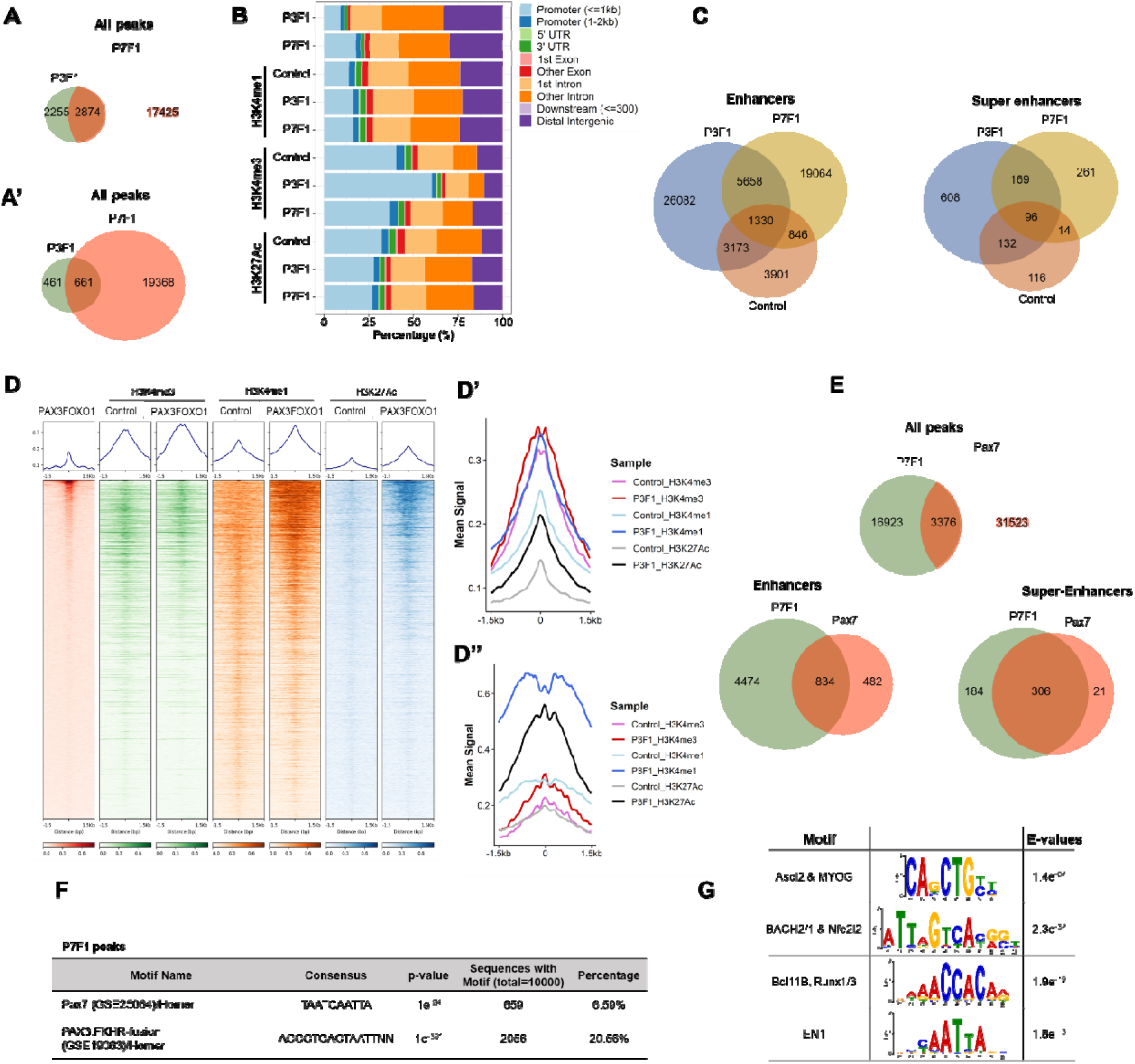
Characterizing genome-wide epigenetic changes in iPSC-MP^PAX3/7-FOXO^ models. **A)** Venn diagram plot depicting overlap of genome-wide binding sites between PAX3-FOXO1 and PAX7-FOXO1 fusion transcription factors in iPSC-MP^PAX3/7-FOXO^ cells. **B)** A composite bar graph depicting percent occupancy of PAX3-FOXO1, PAX7- FOXO1 and histone marks (H3K27Ac, H3K4me1, H3K4me3) at promoter, exonic, intronic and intergenic genomic regions in fusion negative control cells and iPSC- MP^PAX3/7-FOXO1^ cells. **C)** Overlap between enhancers and SE in iPSC-MP^PAX3-FOXO1^ and iPSC-MP^PAX7-FOXO1^ cells. **D)** Composite heatmap of histone modifications signal centered on PAX3-FOXO1 peaks and a window spanning -1.5 to 1.5 kb relative to peaks in iPSC-MP^PAX3-FOXO1^ cells shows enrichment of histone modifications as compared to the same regions in control iPSC-MP cells. Composite mean signal intensity plot for histone marks associated with PAX3-FOXO1 binding peaks from RH4 cells **(D’)** or iPSC-MP^PAX3-FOXO1^ **(D”)** genome-wide. **E)** Venn diagrams depicting overlaps between PAX7 and PAX7-FOXO1 binding across all binding sites and annotated enhancers and super-enhancers (SE) clusters genome-wide. **F)** HOMER- based motif analysis reveals enrichment of PAX7 and PAX3-FOXO1 motifs in PAX7- FOXO1 occupancy sites genome-wide. **G)** Motif analysis of PAX7-FOXO1 peaks genome-wide in iPSC-MP^PAX7-FOXO1^ cells.

**Figure S3.**
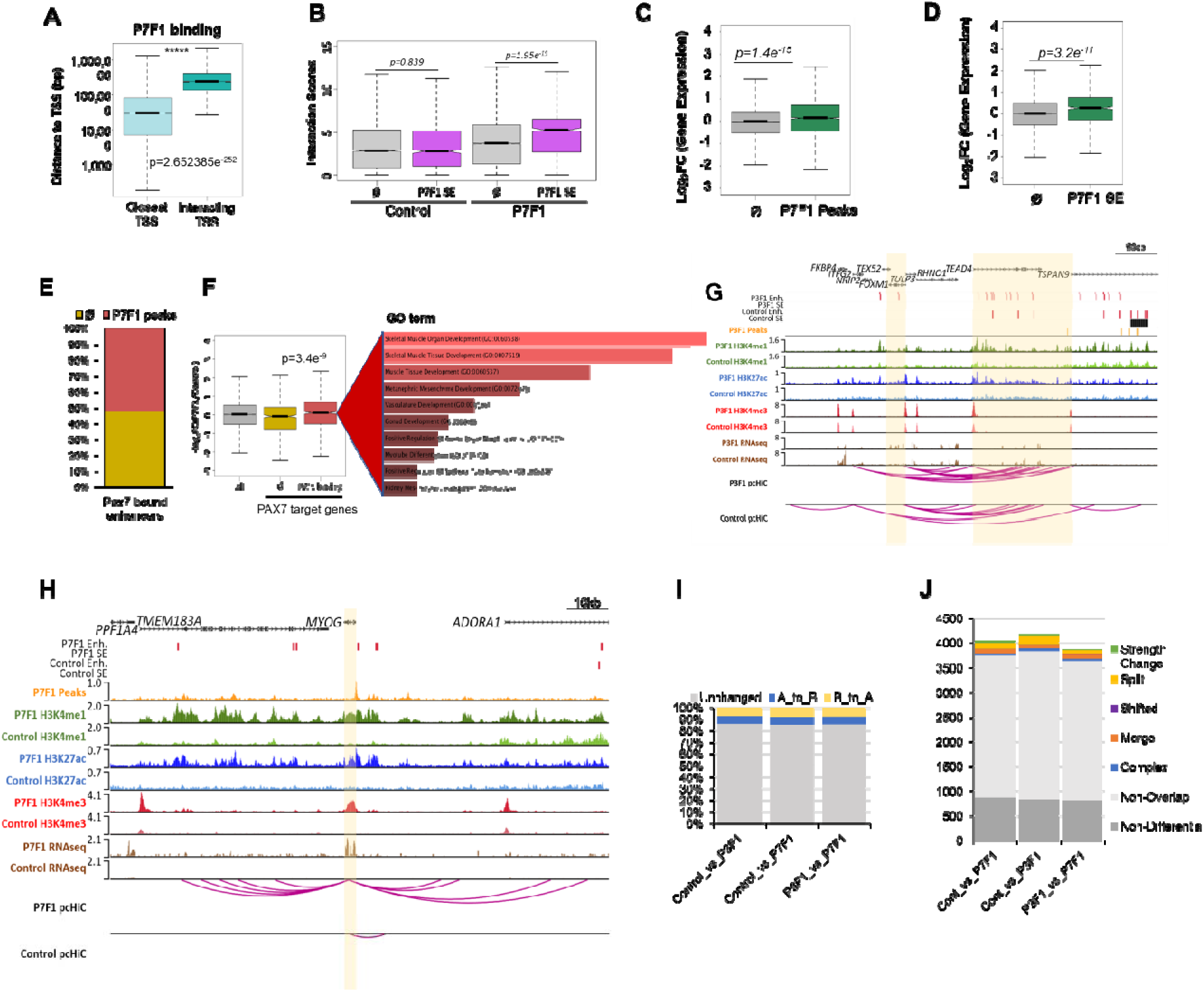
Oncogenic TFs establishes long-range promoter-anchored enhancer interaction loops to regulate its target genes. **A)** Boxplot comparing genomic distance (in log scale) of PAX7-FOXO1 binding sites to its nearest transcription start site (TSS) versus TSS regulated by long-range P-E interaction networks (calculated from pCHi-C data). n = 20299 for closest TSS, n = 2593 for interacting TSS; error bar, SEM; Welch’s t-test. **B)** Boxplot of interaction scores in fusion negative control and iPSC-MP^PAX7-FOXO1^ cells for P-E interactions mediated by PAX7-FOXO1 binding events compared to interactions mediated in absence of PAX7-FOXO1 (Φ) in annotated super-enhancers (SE) regions. n = 8091, 877 for control, n = 8091, 877 for P7F1; error bar, SEM; Welch’s t-test. **C)** Boxplot of normalized log_2_ fold-changes in gene expression (iPSC-MP^PAX7-FOXO1^ vs Control) for P-E interactions mediated by PAX7-FOXO1 binding events compared to interactions observed in the absence of PAX7-FOXO1 (Φ). n = 6375 for Φ, n = 2593 for P7F1; error bar, SEM; Welch’s t-test. **D)** Boxplot of normalized log_2_-fold changes in gene expression (iPSC-MP^PAX7-FOXO1^ vs Control) for P-E interactions mediated by PAX7-FOXO1 binding to super enhancer (SE) regions and other (non-SE) regions. n = 8091 for Φ, n = 877 for P7F1; error bar, SEM; Welch’s t-test. **E)** Composite bar graph showing percentage of PAX7-FOXO1 occupancy sites at PAX7-bound enhancers. **F)** Boxplot of normalized log2 fold change in gene expression (iPSC-MP^PAX7-FOXO1^ vs Control) of genome-wide PAX7 target genes in presence or absence (Φ) of PAX7-FOXO1 occupancy. Gene ontology (GO) terms for biological process categories enriched by PAX7-FOXO1 occupancy. n = 8968 for all genes, n = 657 for Φ, n = 719 for P7F1; error bar, SEM; Welch’s t-test. **G)** Genome browser tracks encompassing *FOXM1* locus (promoter is highlighted in yellow), showing CUT&RUN-seq (H3K27Ac, H3K4me1, H3K4me3), RNA-seq, PAX3-FOXO1 binding sites, pCHiC probes, and high-confidence pCHiC interactions (magenta arcs) in control iPSC-MP and iPSC-MP^PAX3-FOXO1^ cells after Dox removal. Super-enhancers are only detected in iPSC-MP cells without Dox. **H)** Genome browser tracks encompassing *MYOG* locus (promoter is highlighted in yellow), showing CUT&RUN-seq (H3K27Ac, H3K4me1, H3K4me3), RNA-seq, PAX7-FOXO1 binding sites, pCHiC probes, and high-confidence pCHi-C interactions (magenta arcs) in fusion negative iPSC-MP and iPSC-MP^PAX7-FOXO1^ cells, after doxycycline removal. **I)** Percentage of changes in A/B compartments in fusion-negative controls (iPSC-MP) and iPSC-MP^PAX3/7-FOXO1^ cells. Approximately 85% of the compartments remained unchanged by the transcriptional activities of PAX3-FOXO1 and PAX7-FOXO1 TFs, whereas ∼7% change from A to B (silencing), and 8% change from B to A (activation). **J)** Genome-wide comparison and annotation of changes associated with topologically associated domain (TADs) boundaries in fusion-negative controls (iPSC-MP) and iPSC-MP^PAX3/7-FOXO1^ cells.

**Figure S4.**
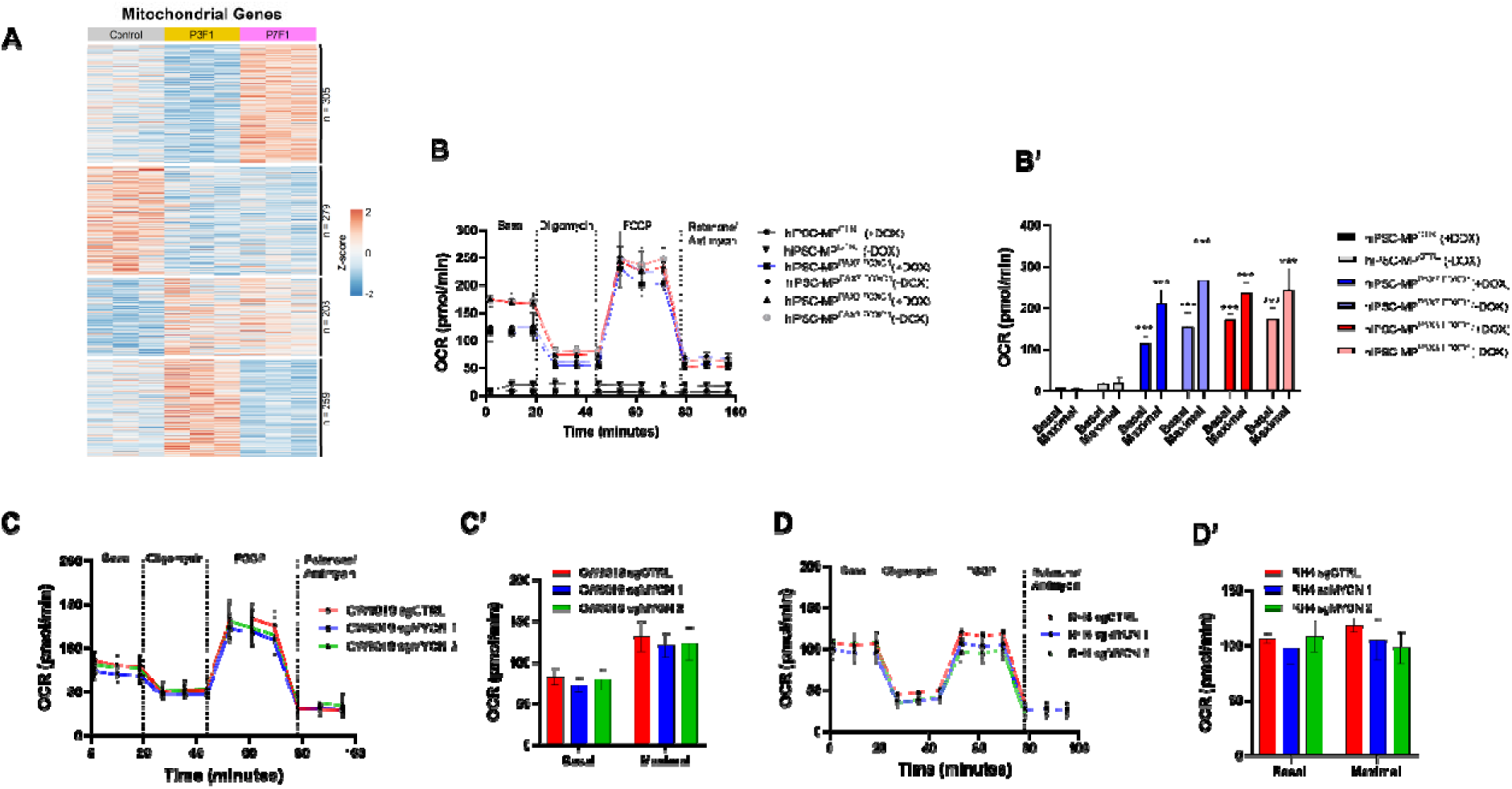
High OXPHOS activity in rhabdomyosarcoma. **A)** Heatmap showing transcriptomic profile of mitochondria-related genes in iPSC-MP cells expressing PAX3-FOXO1 (P3F1) and PAX7-FOXO1 (P7F1) TF and fusion negative control cells. **B)** Seahorse Assay profiles for iPSC-MP cells expressing PAX3-FOXO1 (P3F1) and PAX7-FOXO1 (P7F1) TF and fusion negative control cells, as indicated, using Cell Mito Stress assay. **B’)** Quantification of basal and maximal oxygen consumption rates (OCR) shown in panel B. n = 3 per group; error bar, SEM; **p < 0.01, ***p < 0.001, ****p < 0.0001, ANOVA, Dunnett’s test. Seahorse Assay profiles in RH4 **(C)** and CW9019 **(D)** cells expressing control guides and MYCN-specific gRNAs. **(C’, D’)** Quantification of basal and maximal oxygen consumption rates (OCR) shown in panels C and D. n = 3 per group; error bar, SEM.

**Figure S5.**
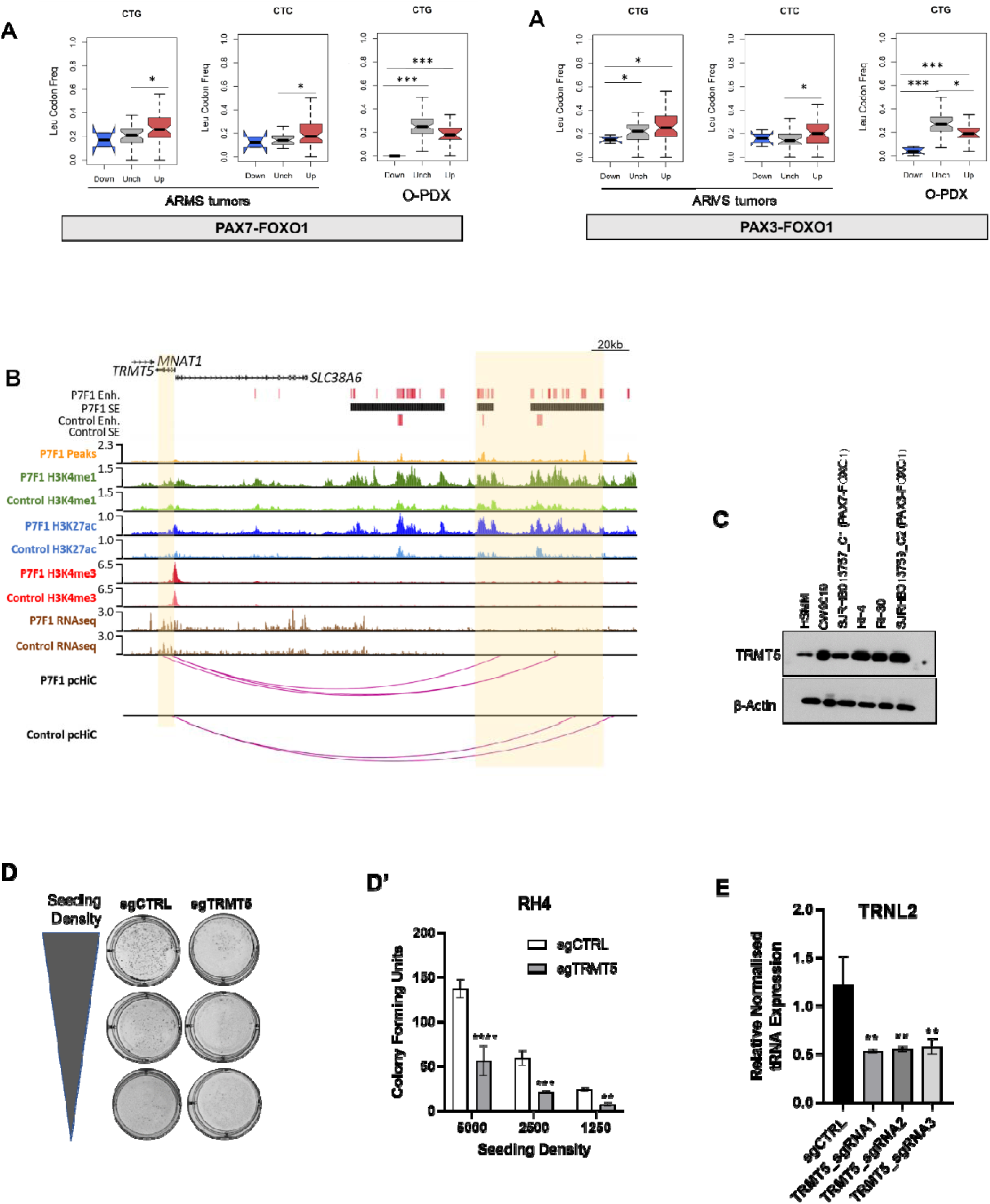
Depletion of *TRMT5* deregulates mitochondrial tRNAs in alveolar rhabdomyosarcoma. **A-A’)** Quantitative estimation of leucine amino acid specific codon bias frequency in **A)** PAX7-FOXO1 ARMS primary tumors and O-PDX samples and **A’)** PAX3-FOXO1 ARMS primary tumors and O-PDX samples. n = 26 per group for ARMS tumors, n = 4 per group for O-PDXs; error bar, SEM; *p < 0.05, ***p < 0.001, ANOVA, Dunnett’s test. **B)** Genome browser tracks encompassing *TRMT5* locus (promoter is highlighted in yellow), showing CUT&RUN (H3K27Ac, H3K4me1, H3K4me3), RNA-seq, PAX7-FOXO1 binding sites, pCHiC probes, and high-confidence pCHiC interactions (magenta arcs) in control iPSC-MP and iPSC-MP^PAX7-FOXO1^ cells after Dox removal. Super-enhancers are only detected in iPSC-MP^PAX7-FOXO1^ cells without Dox. **C)** Western blot images of TRMT5 and β-Actin in HSMM, CW9019, SJRHB013757_C1, RH4, RH30, and SJRHB013759_C2 samples. **D-D’)** Representative images showing colony forming units (CFU) in RH4 stably expressing control and TRMT5-specific sgRNAs seeded at different densities and their respective quantification. n = 3 per group; error bar, SEM; **p < 0.01, ***p < 0.001, ****p < 0.0001, ANOVA, Dunnett’s test. **E)** Transcript levels of *TRNL2* after CRISPRi silencing in RH4 cells using three independent sgRNAs targeting *TRMT5* promoter. n = 3 per group; error bar, SEM; **p < 0.01, ANOVA, Dunnett’s test.

**Figure S6.**
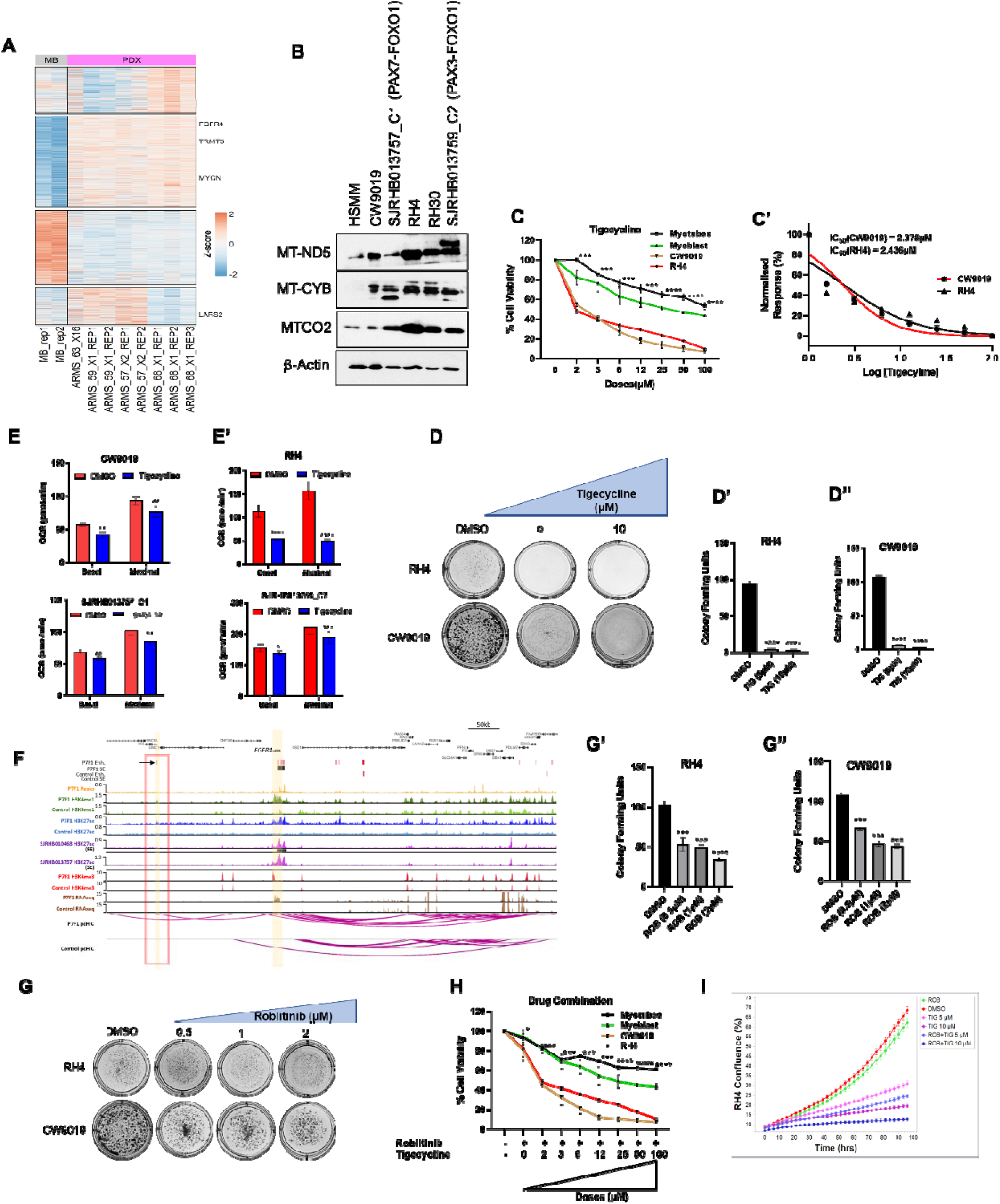
Pharmacological targeting of mitochondrial metabolism and FGFR4 in rhabdomyosarcoma. **A)** Heatmap of protein levels in human myoblast, PAX3-FOXO1 (59_X1, 63_X16) and PAX7-FOXO1 (57_X2, 68_X1) O-PDX samples. **B)** Western blot images of MT-ND5, MT-CYB, MTCO2 and β-Actin in HSMM, CW9019, SJRHB013757_C1, RH4, RH30, and SJRHB013759_C2 samples. **C)** Cell viability curve of human myotubes, myoblast, CW9019 and RH4 cells in response to Tigecycline treatment at various doses. n = 3 per group; error bar, SEM; ***p < 0.001, ****p < 0.0001, ANOVA, Dunnett’s test. **C’)** Derivation of IC50 values for RH4 and CW9019 cells treated with various doses of Tigecycline. **D-D’’)** Representative images showing colony forming units (CFU) in RH4 and CW9019 cells treated with Tigecycline (5 μM, 10 μM) and their respective quantification **(D’, D’’)**. n = 3 per group; error bar, SEM; ****p < 0.0001, ANOVA, Dunnett’s test. Estimation of oxygen consumption rate (OCR) in **E)** CW9019 and SJRHB013757_C1 2D-adapted PDX model **E’)** RH4 and SJRHB013759_C2 2D-adapted PDX models treated with Tigecycline (10 μM). n = 3 per group; error bar, SEM; *p < 0.05, **p < 0.01, ***p < 0.001, ****p < 0.0001, ANOVA, Dunnett’s test. **F)** Genome browser tracks encompassing *FGFR4* locus (promoter highlighted in yellow), showing CUT&RUN (H3K27Ac, H3K4me1, H3K4me3), RNA-seq, PAX7-FOXO1 binding sites, pCHiC probes, and high-confidence pCHiC interactions (magenta arcs) in control iPSC-MP and iPSC-MP^PAX7-FOXO1^ cells after Dox removal and PAX7-FOXO1 positive O-PDXs (SJRHB0104681, SJRHB013757)^6^. Indicated super enhancers (black bar) and promoter-anchored upstream enhancer (∼180kb, arrow signs indicated within red rectangle) are only detected in iPSC-MP^PAX7-FOXO1^ cells without Dox. **G)** Representative images showing colony forming units (CFU) in RH4 and CW9019 cells treated with Roblitinib at indicated dosages. (**G’, G”)** Quantification of colony forming units (CFU) in panel G. n = 3 per group; error bar, SEM; ***p < 0.001, ****p < 0.0001, ANOVA, Dunnett’s test. **H)** Cell viability curves for human myotubes, myoblasts, CW9019, and RH4 cells in response to Tigecycline and Roblitinib treatments at indicated doses. n = 3 per group; error bar, SEM; *p < 0.05, ***p < 0.001, ****p < 0.0001, ANOVA, Dunnett’s test. **I)** Cell confluency curves for RH4 cells in response to Tigecycline (5μM, 10μM), Roblitinib (1μM) (ROB+TIG) combination treatment.

